# Meiosis-specific ZFP541 repressor complex promotes meiotic prophase exit during spermatogenesis

**DOI:** 10.1101/2021.01.15.426901

**Authors:** Yuki Horisawa-Takada, Chisato Kodera, Kazumasa Takemoto, Akihiko Sakashita, Kenichi Horisawa, Ryo Maeda, Shingo Usuki, Sayoko Fujimura, Naoki Tani, Kumi Matsuura, Ryuki Shimada, Tomohiko Akiyama, Atsushi Suzuki, Hitoshi Niwa, Makoto Tachibana, Takashi Ohba, Hidetaka Katabuchi, Satoshi H. Namekawa, Kimi Araki, Kei-ichiro Ishiguro

## Abstract

During spermatogenesis, meiosis is accompanied by robust alteration in gene expression and chromatin status. However, it remained elusive how meiotic transcriptional program is established to ensure completion of meiotic prophase. Here, we identified a novel protein complex consisting of germ-cell-specific zinc-finger protein ZFP541 and its interactor KCTD19 as the key transcriptional regulator for meiotic prophase exit. Our genetic study showed that ZFP541 and KCTD19 are co-expressed from pachytene onward and play an essential role in the completion of meiotic prophase program in the testis. Furthermore, our ChIP-seq and transcriptome analyses revealed that ZFP541 binds to and suppresses a broad range of genes whose function is associated with biological processes of transcriptional regulation and covalent chromatin modification. The present study demonstrated that germ-cell specific ZFP541-KCTD19 containing complex promotes meiotic prophase exit in males, and triggers reconstruction of the transcription network and chromatin organization leading to post-meiotic development.

## Introduction

Meiosis consists of one round of DNA replication followed by two rounds of chromosome segregation, producing haploid gametes from diploid cells. Meiotic entry is followed by meiotic prophase that corresponds to a prolonged G2 phase in which meiosis-specific chromosomal events such as chromosome axis formation, homolog synapsis and meiotic recombination sequentially occur. (Handel and Schimenti, 2010) (Zickler and Kleckner, 2015) (Page and Hawley, 2004). Completion of meiotic prophase is regulated by sexually dimorphic mechanisms, so the transcription, cell cycle and chromatin status are altered in subsequent developmental program for sperm production and oocyte arrest/maturation. In male germ cells, the completion of pachytene is monitored under several layers of regulation such as pachytene checkpoint and meiotic sex chromosome inactivation (MSCI) (Turner, 2015) (Burgoyne et al., 2009) (Ichijima et al., 2012). Male meiotic prophase is accompanied by robust changes of gene expression programs (Namekawa et al., 2006) (Shima et al., 2004) (Schultz et al., 2003) and epigenetic status (Sin et al., 2015) (Maezawa et al., 2020) (Kota and Feil, 2010) (Sasaki and Matsui, 2008) as well as reorganization of the chromatin structure (Alavattam et al., 2019) (Wang et al., 2019) (Kimmins and Sassone-Corsi, 2005) for post-meiotic development. At pachytene stage of male meiotic prophase, the transcriptional program for postmeiotic stages starts to take place (Sin et al., 2015) (da Cruz et al., 2016) (Ernst et al., 2019). A germline-specific Polycomb protein, SCML2 is at least in part responsible for the suppression of somatic/progenitor genes and the activation of late-spermatogenesis-specific genes in pachytene spermatocytes and round spermatids (Hasegawa et al., 2015) (Maezawa et al., 2018a) (Maezawa et al., 2018b). Thus for spermatocytes, passing through the pachytene stage during meiotic prophase is a critical developmental evet for subsequent spermatid differentiation. However, it remained elusive how meiotic transcriptional program is established to ensure completion of meiotic prophase prior to post-meiotic differentiation.

In this study, we identified a novel protein complex that establishes meiotic transcription program to ensure completion of meiotic prophase. Previously, we identified MEIOSIN that plays an essential role in meiotic initiation both in male and female (Ishiguro et al., 2020). MEIOSIN together with STRA8 (Kojima et al., 2019) acts as a crucial transcription factor that drives meiotic gene activation. In the present study, *Zfp541* gene, that encodes a zinc finger protein, was identified as one of the MEIOSIN/STRA8-target genes. Although ZFP541 was previously identified as a Zinc finger protein, SHIP1, in the spermatocyte UniGene library showing testis-specific expression (Choi et al., 2008), its function is yet to be determined. Here we show that ZFP541 plays a role in promoting pachytene exit in male germ cells. By mass spectrometry (MS) analysis, we showed that ZFP541 is expressed at pachytene and interacts with KCTD19 and HDAC1/2. Disruption of *Zfp541* and *kctd19* led to defects in the completion of meiotic prophase, with severe impact on male fertility. Chromatin binding analysis of ZFP541 combined with transcriptome analysis demonstrated that ZFP541 binds to and represses a broad range of genes whose biological functions are associated with the processes of transcriptional regulation and covalent chromatin modification. In round spermatids, those ZFP541-target genes were repressed with repressive histone marks. The present study suggests that ZFP541-containg complex globally represses the gene expression in meiotic prophase, and triggers the reconstruction of transcription and chromatin organization to proceed with the developmental program for sperm production.

## Results

### ZFP541 is expressed during meiotic prophase in spermatocytes and oocytes

Previously, we identified a germ cell specific transcription factor, MEIOSIN, that directs initiation of meiosis (Ishiguro et al., 2020). Our study demonstrated that MEIOSIN works with STRA8 and activates hundreds of meiotic genes, which are required for meiotic initiation. We identified *Zfp541* as one of the MEIOSIN/STRA8-bound genes during preleptotene, the time of meiotic initiation (Fig 1A). Expression of *Zfp541* was significantly downregulated in RNA-seq analysis of *Meiosin* KO testes at postnatal day 10 (P10) (Ishiguro et al., 2020) which is the time when a cohort of spermatocytes undergo the first wave of meiotic entry. We confirmed this by RT-qPCR analysis demonstrating that *Zfp541* expression level was downregulated in *Meiosin* KO testis at P10 (Fig. 1B). We further examined the expression pattern of the *Zfp541* in different mouse tissues by RT-PCR analysis. *Zfp541* gene showed a specific expression in adult and embryonic testes, and in embryonic ovary, but not in other adult organs that we examined (Fig. 1C, D), suggesting that *Zfp541* is a germ-cell-specific factor both in male and female, which is partly consistent with previous study (Choi et al., 2008).

**Figure 1.**
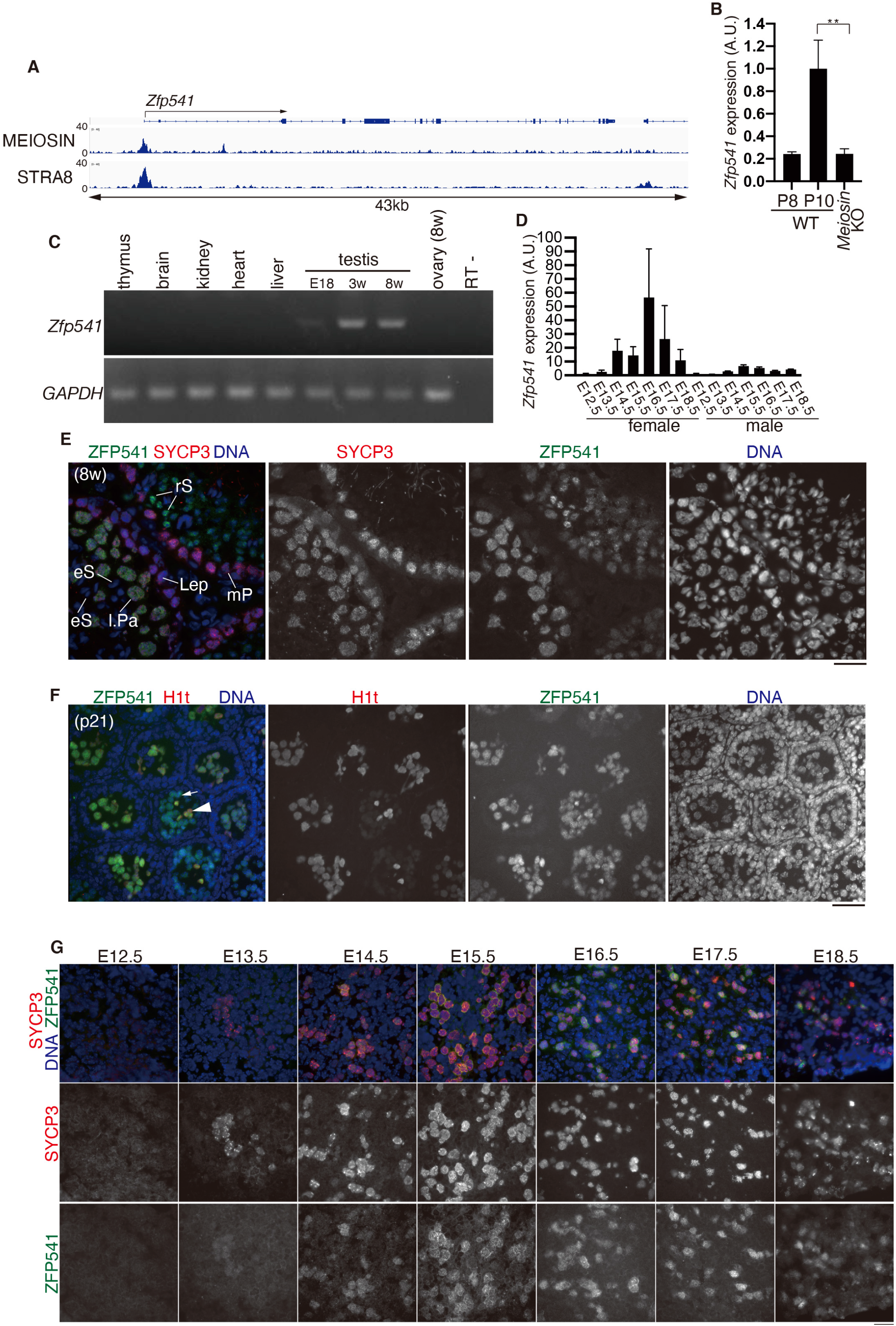
*Zfp541* was identified as a MEIOSIN-STRA8 target gene. **(A)** Genomic view of MEIOSIN and STRA8 binding peaks over the *Zfp541* locus. Genomic coordinates derived from Ensembl. **(B)** The expression of *Zfp541* in WT (P8 and P10) and *Meiosin* KO testes was examined by RT-qPCR. Three animals for each genotype were used. The graph shows the expression level of *Zfp541* normalized by that of *GAPDH* with SD. Expression level of *Zfp541* in P10 WT was set to 1. Statistical significance is shown by *p*-value (t-test).**: p < 0.01. **(C)** The tissue-specific expression pattern of *Zfp541* was examined by RT-PCR. Testis RNA was obtained from embryonic day 18 (E18), 3-weeks old (3w) and 8-weeks old (8w) male mice. Ovary RNA was obtained from adult 8-weeks old (8w) female mice. RT-indicates control PCR without reverse transcription. **(D)** The expression patterns of *Zfp541* in the embryonic ovary and testis were examined by RT-qPCR. **(E)**Seminiferous tubule sections in WT testis (8-weeks old) were immunostained as indicated. Lep: leptotene, mP: mid pachytene, lPa: late pachytene spermatocyte, rS: round spermatid, eS: elongated spermatid. **(F)** Seminiferous tubule sections in WT testis (P21) were immunostained as indicated. Arrow and arrowhead indicate H1t-negative and H1t-positive pachytene spermatocyte, respectively. **(G)** Embryonic ovary sections were stained as indicated. Scale bars: 25 μm.

It has been suggested that ZFP541 possesses putative DNA-binding domains, C2H2 type zinc-finger and ELM2-SANT domains (Choi et al., 2008). However, its biological function has remained elusive. To determine the meiotic stage/cell type-specific expression of ZFP541, the seminiferous tubules of the WT testes (8-weeks old) were immunostained with specific antibodies against ZFP541 along with SYCP3 (a component of meiotic axial elements) (Fig. 1E). ZFP541 appeared in the nuclei of the spermatocytes from pachytene onward, and in round spermatids (Fig. 1E). However, it was not observed in spermatocytes before the stages of pachytene, elongated spermatids, or spermatogonia. Testis-specific histone H1t is a marker of spermatocytes later than mid pachytene (Cobb et al., 1999) (Drabent et al., 1996). Immunostaining of seminiferous tubules by testis-specific histone H1t indicated that localization of the ZFP541 protein into the nuclei started at H1t negative early pachytene stage (Fig. 1F). Although the expression of *Zfp541* mRNA was upregulated upon meiotic entry, immunostaining of ZFP541 protein detected no more than background levels in spermatocytes before pachytene stage (Fig. 1F), suggesting that expression of ZFP541 may be post-transcriptionally regulated upon the entry into meiotic prophase I. In females, ZFP541 was expressed in the SYCP3 positive oocytes in embryonic ovary during E13.5 −18.5, the stage when oocytes progressed the meiotic prophase (Fig. 1G). Thus, ZFP541 was expressed in meiotic prophase both in male and female.

### *Zfp541* knockout in mice results in male infertility

In order to address the role of ZFP541, we deleted Exon1-Exon13 of the *Zfp541* loci in C57BL/6 fertilized eggs by CRISPR/Cas9 mediated genome editing (Fig. 2A). Immunoblotting of the extract from *Zfp541* KO testis showed that ZFP541 protein was absent (Fig. 2B), which was further confirmed by diminished immunolocalization of ZFP541 in the seminiferous tubules of *Zfp541* KO (Fig. 2F), indicating that the targeted *Zfp541* allele was null. Although *Zfp541* KO mice developed normally, defects in male reproductive organs were evident with smaller-than-normal testes (Fig. 2C). Histological analysis revealed that post-meiotic spermatids and spermatozoa were absent in the *Zfp541* KO seminiferous tubules (Fig. 2D, F). Accordingly, sperm was absent in *Zfp541* KO caudal epididymis (Fig. 2E). Thus, the later stage of spermatogenesis was severely abolished in *Zfp541* KO seminiferous tubules, resulting in male infertility. In contrast to male, *Zfp541* KO females exhibited seemingly normal fertility with no apparent defects in adult ovaries (Fig. S1A, B). Therefore, these results suggested that requirement of ZFP541 is sexually different.

**Figure 2.**
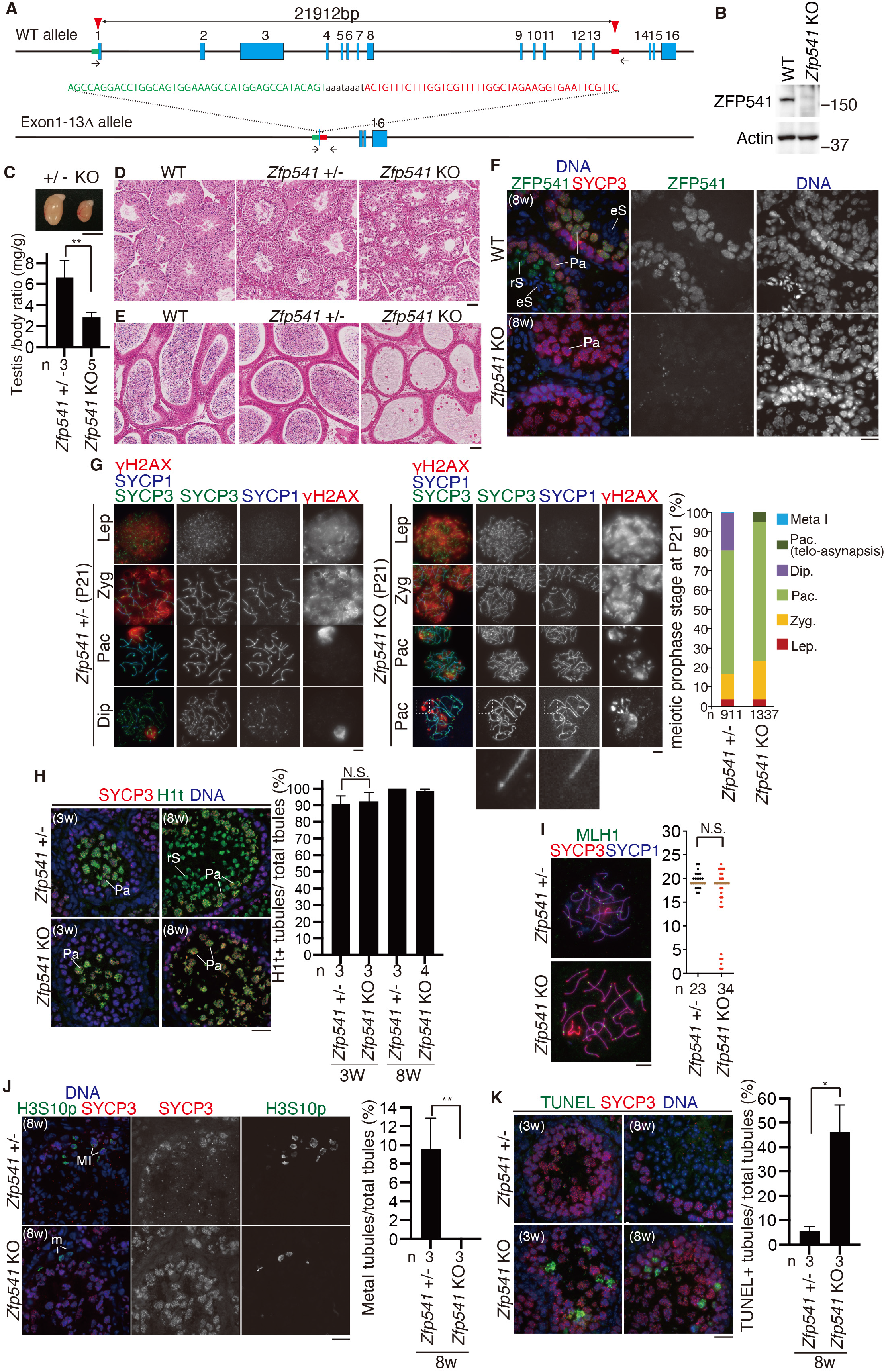
*Zfp541* knockout mice are infertile in male. **(A)** The allele with targeted deletion of Exon1-13 in *Zfp541* gene was generated by the introduction of CAS9, the synthetic gRNAs designed to target Exon1 and the downstream of Exon13 (arrowheads), and ssODN into C57BL/6 fertilized eggs. Arrows: PCR primers for genotyping. **(B)** Immunoblot analysis of testis extracts prepared from mice with the indicated genotypes (P21). **(C)** Testes from *Zfp541* +/− and *Zfp541* KO (8-weeks old). Scale bar: 5 mm. Testis/body-weight ratio (mg/g) of *Zfp541* +/− and *Zfp541* KO mice (8-weeks old) is shown below (bar graph with SD). n: the number of animals examined. Statistical significance is shown by *p*-values (paired t-test). **: *p* <0.001 **(D)** Hematoxylin and eosin staining of the sections from WT, *Zfp541* +/− and *Zfp541* KO testes (8-weeks old). Scale bar: 50 μm. **(E)** Hematoxylin and eosin staining of the sections from WT, *Zfp541* +/− and *Zfp541* KO epididymis (8-weeks old). Scale bar: 50 μm. **(F)** Seminiferous tubule sections (8-weeks old) were stained for SYCP3, ZFP541 and DAPI. Scale bars: 15 μm. Pa: pachytene spermatocyte, rS: round spermatid, eS: elongated spermatid. **(G)** Chromosome spreads of *Zfp541* +/− and *Zfp541* KO spermatocytes at postnatal day 21 (P21) were immunostained as indicated. Scale bar: 5 μm. Enlarged images are shown to highlight asynapsed chromosomes in pachytene-like cell. Lep: leptotene, Zyg: zygotene, Pac: pachytene, Dip: diplotene. Quantification of meiotic prophase stage spermatocytes per total SYCP3+ spermatocytes in *Zfp541* +/− and *Zfp541* KO mice is shown on the right. n: the number of cells examined. **(H)** Seminiferous tubule sections (3-weeks and 8-weeks old) were stained for SYCP3, H1t and DAPI. Scale bars: 25 μm. Shown on the right is the quantification of the seminiferous tubules that have H1t+/SYCP3+ cells per the seminiferous tubules that have SYCP3+ spermatocyte cells in *Zfp541* +/− and *Zfp541* KO mice (bar graph with SD). n: the number of animals examined for each genotype. Statistical significance is shown (paired t-test). N.S.: Statistically not significant. **(I)** Chromosome spreads of *Zfp541* +/− and *Zfp541* KO spermatocytes were stained for MLH1, SYCP3 and SYCP1. Scale bars: 5 μm. The number of MLH1 foci is shown in the scatter plot with median (right). Statistical significance is shown (Mann-Whitney U-test). n: number of spermatocytes examined. N.S.: Statistically not significant. **(J)** Seminiferous tubule sections (8-weeks old) were stained for SYCP3, H3S10P and DAPI. Scale bars: 25 μm. M I: Metaphase I spermatocyte, m: Metaphase of mitotic spermatogonia. Shown on the right is the quantification of the seminiferous tubules that have H3S10p+ /SYCP3+ Metaphase I cells per the seminiferous tubules that have SYCP3+ spermatocyte cells in *Zfp541* +/− and *Zfp541* KO testes (bar graph with SD). n: the number of animals examined for each genotype. Statistical significance is shown by *p*-value (t-test). **: p < 0.01. Scale bar: 25 μm. **(K)** Seminiferous tubule sections from 3-weeks and 8-weeks old mice were subjected to TUNEL assay with immunostaining for SYCP3. Shown on the right is the quantification of the seminiferous tubules that have TUNEL+ cells per total tubules in *Zfp541* +/− (8w; n=3) and *Zfp541* KO (8w; n=3) testes (bar graph with SD). Statistical significance is shown by *p*-values (paired t-test). *: *p* <0.05. Scale bar: 25 μm.

### ZFP541 is required for pachytene exit in male

To further investigate at which stage the primary defect appeared in the *Zfp541* KO, we analyzed the progression of spermatogenesis by immunostaining. Immunostaining analysis with antibodies against SYCP3 (a component of meiotic chromosome axis) along with SYCP1 and γH2AX (a marker of DNA damage response for meiotic recombination and XY body), demonstrated that spermatocytes initiated double strand breaks (DSBs) for meiotic recombination and underwent homologous chromosome synapsis in juvenile *Zfp541* KO males (P21) as in age-matched WT (Fig. 2G). However, spermatocytes later than pachytene did not appear in *Zfp541* KO mice at P21 (Fig. 2G), whereas the first wave of spermatogenesis passed through the meiotic prophase in the age-matched control. Notably, γH2AX still remained along synapsed autosomes despite the completion of homolog synapsis, and 6.9% of pachytene nuclei exhibited homolog chromosomes partially lacking SYCP1 assembly at the telomeric region in *Zfp541* KO at P21 (Fig. 2G). This suggested that DNA damage response still remained active along synapsed homologs in *Zfp541* KO. Close inspection of the seminiferous tubules indicated that *Zfp541* KO spermatocytes reached to at least late pachytene as indicated by the presence of H1t staining (Fig. 2H). The number of MLH1 foci (a marker of crossover recombination) in *Zfp541* KO pachytene spermatocytes was comparable to age-matched control (Fig. 2I), suggesting that crossover recombination was complete in *Zfp541* KO pachytene spermatocytes, even though DNA damage response was still active. γH2AX signal was observed on the XY body in *Zfp541* KO spermatocytes (Fig. 2G), suggesting that silencing of XY chromosomes took place in *Zfp541* KO spermatocytes. These observations suggested that the progression of meiotic prophase beyond pachytene was compromised in *Zfp541* KO. Accordingly, *Zfp541* KO spermatocytes failed to reach metaphase I stage as indicated by immunostaining of Histone H3 Ser10 phosphorylation (H3S10P) and centromeric SYCP3 (Fig. 2J). Hence, the primary defect had already occurred before the completion of meiotic prophase rather than during spermatid development. Notably, TUNEL positive cells were observed in the stage X-XI seminiferous tubules that contained late pachytene spermatocytes (Fig. 2K). The same phenotype persisted through adulthood in *Zfp541* KO testes, because higher number of TUNEL positive seminiferous tubules (~46% in total tubules) were observed in *Zfp541* KO testes at 8-weeks old (Fig. 2K). These observations suggested that *Zfp541* KO spermatocytes failed to exit pachytene and were consequently eliminated by apoptosis. These results indicate ZFP541 is required for exiting pachytene stage in spermatogenesis.

### *Kctd19* knockout in mice results in male infertility

In order to elucidate the function of ZFP541, the interacting factors of ZFP541 were screened by immunoprecipitation (IP) followed by mass spectrometry (MS) analysis. Our ZFP541 IP-MS analysis demonstrated that ZFP541 associated with histone deacetylases HDAC1, HDAC2, POZ/BTB domain containing KCTD19 and terminal deoxynucleotidyltransferase interacting factor 1 (TDIF1) in the chromatin-bound fractions of testes extracts (Fig. 3A, Fig. S2), which was consistent with previous study (Choi et al., 2008). Similar results were reproducibly obtained by MS analysis of ZFP541-IP from the chromatin-unbound fraction and the MNase-released nucleosome fractions of testes extracts (Fig. S2). Furthermore, reciprocal IP by KCTD19 antibody indicated that ZFP541, HDAC1, HDAC2 and TDIF1 were co-immunoprecipitated with KCTD19 (Fig. 3A, Fig. S3). These results suggest that those factors form a complex and may play a role in transcriptional repression through histone deacetylases. It was previously shown that TDIF1 (Bantscheff et al., 2011) (Zhang et al., 2018) (Sawai et al., 2018) associated with histone deacetylases and ELM2-SANT domain containing MIDEAS/ELMSAN1, and knockdown of TDIF1 led to mitotic chromosome misalignment (Turnbull et al., 2020). However, the exact function of KCTD19 has remained elusive. RT-PCR analyses demonstrated that *Kctd19* gene showed a specific expression in adult testes, and not in other adult organs we examined (Fig.3B, Fig. S4A), suggesting that *Kctd19* is a germ-cell-specific factor. Similar to the expression pattern of ZFP541, KCTD19 appeared in the nuclei of the spermatocytes from pachytene onward, and in round spermatids (Fig. 3C, D). Accordingly, KCTD19 was co-expressed with its binding partner ZFP541 in testis (Fig. 3E). In females, although *Kctd19* mRNA was detected in embryonic ovaries by RT-PCR (Fig. S4A), protein expression of KCTD19 in embryonic ovaries was under detection limit by immunostaining (Fig. S4B).

**Figure 3.**
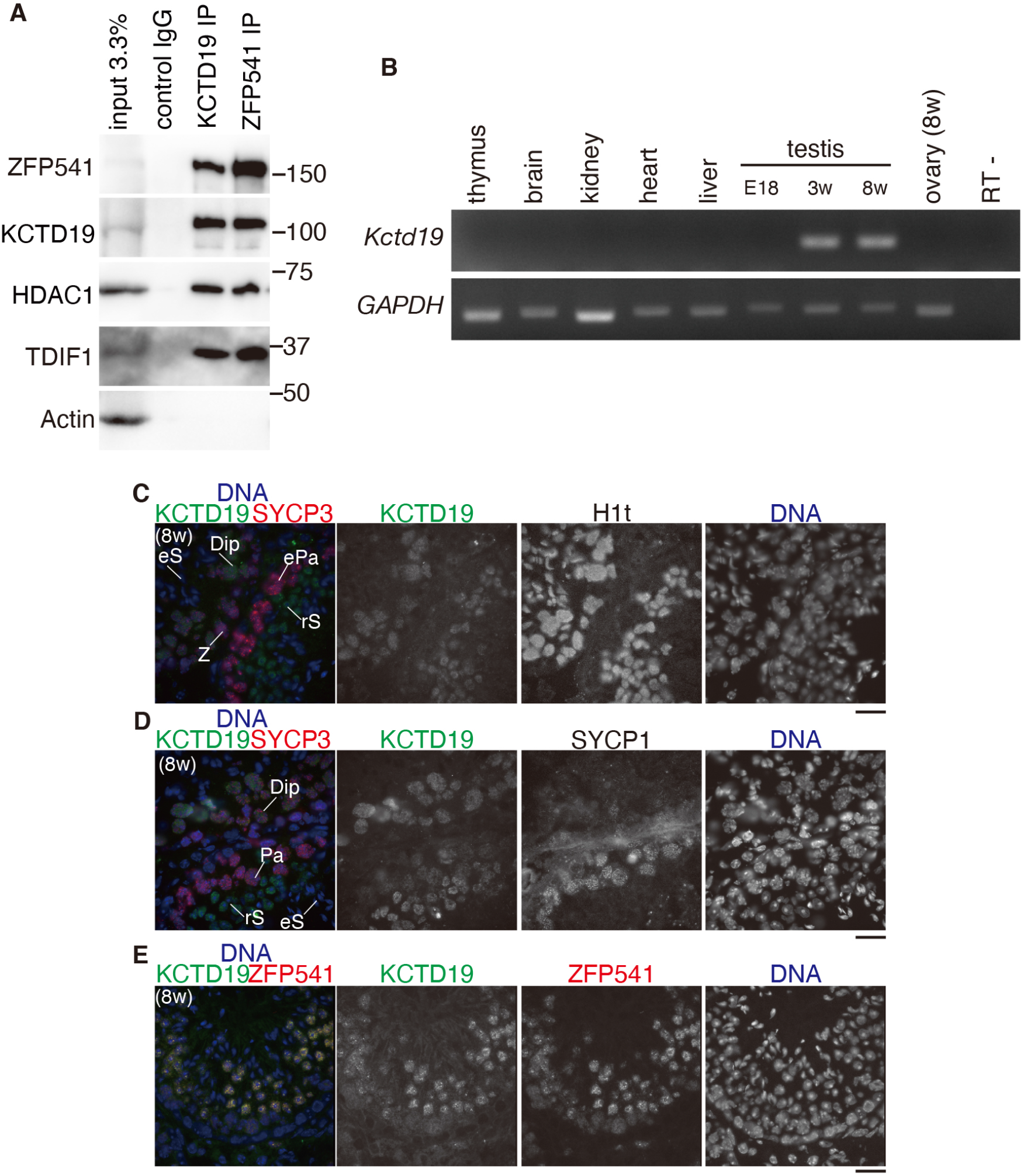
KCTD19 was identified as a ZFP541 interacting factor in the testis. **(A)** Immunoblot showing the immunoprecipitates of ZFP541 and KCTD19 from chromatin extracts of WT mouse testes. **(B)** The tissue-specific expression pattern of *Kctd19* was examined using RT-PCR. Testis RNA was obtained from embryonic day 18 (E18), 3-weeks old (3w) and 8-weeks old (8w) male mice. Ovary RNA was obtained from adult 8-weeks old (8w) female mice. **(C-E)** Seminiferous tubule sections in WT testis (8-weeks old) were immunostained as indicated. Z: zygotene, Pa: pachytene, ePa: early pachytene, Dip: diplotene, rS: round spermatid, eS: elongated spermatid. Scale bars: 25 μm.

In order to address the role of KCTD19, we deleted Exon3-Intron12 of the *Kctd19* loci in C57BL/6 fertilized eggs by CRISPR/Cas9 mediated genome editing (Fig. 4A). In *Kctd19* KO testis, ZFP541 protein was expressed, and in *Zfp541* KO testes KCTD19 was expressed (Fig. 4B), implying that the expressions of ZFP541 and KCTD19 are independent of each other. Notably, whereas ZFP541 localized in the nuclei of the *Kctd19* KO seminiferous tubules, KCTD19 was not detected in the nuclei of the *Zfp541* KO seminiferous tubules despite the expression of KCTD19 in *Zfp541* KO testes, suggesting that nuclear localization of KCTD19 depends on the presence of ZFP541 (Fig. 4C). Similar to *Zfp541* KO mice, whereas the *Kctd19* KO females exhibited seemingly normal fertility with no apparent defect in adult ovaries (Fig. S4C, D), *Kctd19* KO males exhibited smaller-than-normal testes (Fig. 4D). Indeed, histological analyses of *Kctd19* KO males revealed impaired spermatogenesis shown by post-meiotic spermatids and spermatozoa being absent in the seminiferous tubules and the caudal epididymis (Fig. 4E, F). These observations suggested that KCTD19 is required for normal progression of spermatogenesis.

**Figure 4.**
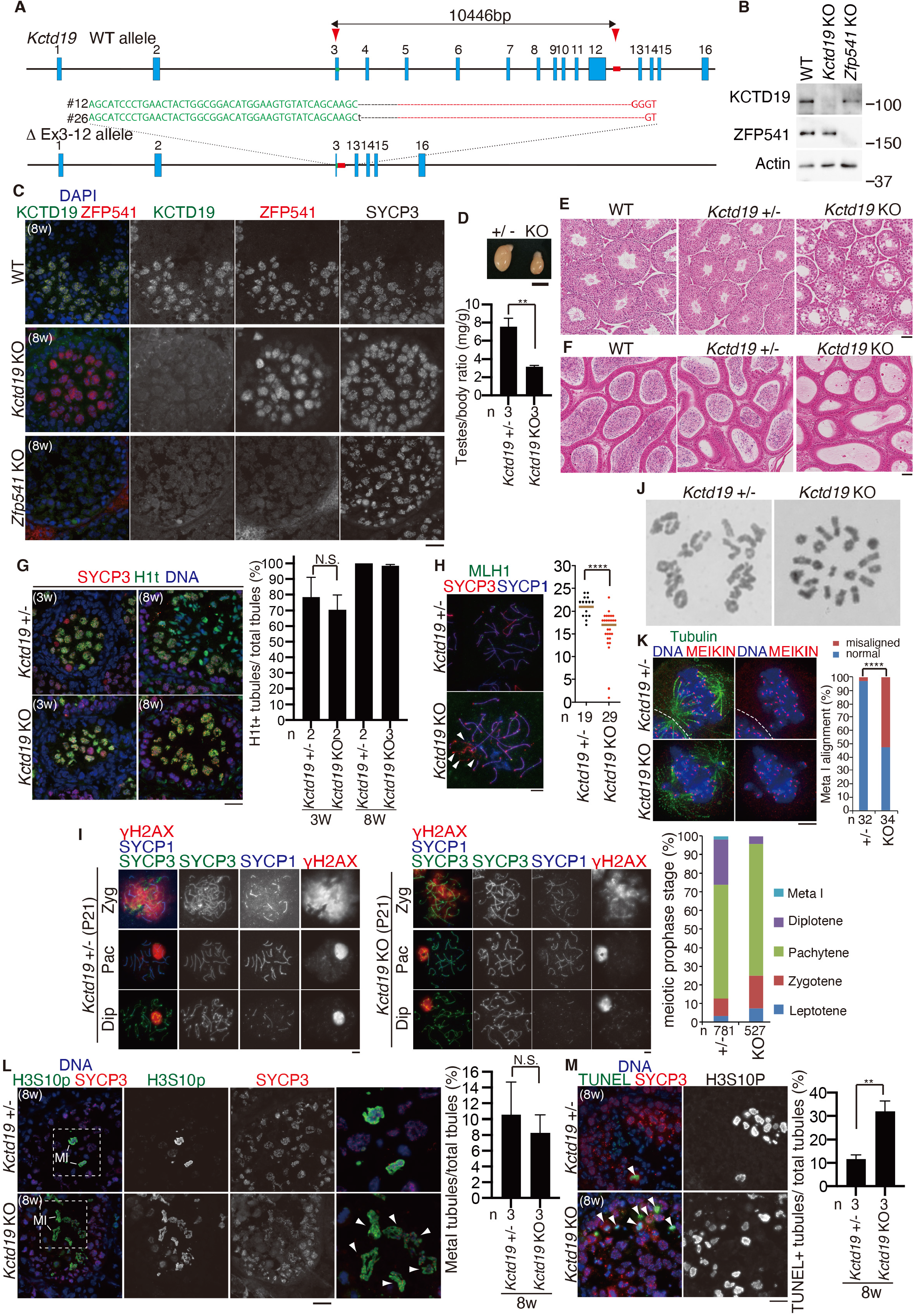
KCTD19 is required for completion of meiosis I. **(A)** The allele with targeted deletion of Exon3-12 in *Kctd19* gene were generated by the introduction of CAS9, the synthetic gRNAs designed to target Exon 3 and the downstream of Exon12 (red arrowheads), and ssODN into C57BL/6 fertilized eggs. Two lines of KO mice were established. Line #12 of *Kctd19* KO mice was used in most of the experiments, unless otherwise stated. **(B)** Immunoblot analysis of whole testis extracts prepared from mice with the indicated genotypes (P21). **(C)** Seminiferous tubule sections from WT, *Kctd19* KO and *Zfp541* KO (8-weeks old) were immunostained as indicated. Scale bar: 25 μm. **(D)** Testes from *Kctd19* +/− and *Kctd19* KO (8-weeks old). Scale bar: 5 mm. Testis/body-weight ratio (mg/g) of *Kctd19* +/− and *Kctd19* KO mice (8-weeks old) is shown below (bar graph with SD). n: number of animals examined. Statistical significance is shown by *p*-values (paired t-test). **: *p* <0.01 **(E)** Hematoxylin and eosin staining of the sections from WT, *Kctd19* +/− and *Kctd19* KO testes (8-weeks old). Scale bar: 50 μm. **(F)** Hematoxylin and eosin staining of the sections from WT, *Kctd19* +/− and *Kctd19* KO epididymis (8-weeks old). Scale bar: 50 μm. **(G)** Seminiferous tubule sections (3-weeks and 8-weeks old) were immunostained as indicated. Scale bar: 25 μm. Shown on the right is the quantification of the seminiferous tubules that have H1t+/SYCP3+ cells per the seminiferous tubules that have SYCP3+ spermatocyte cells in *Kctd19* +/− and *Kctd19* KO mice (bar graph with SD). n: the number of animals examined for each genotype. Statistical significance is shown (paired t-test). N.S.: Statistically not significant. **(H)** Chromosome spreads of *Kctd19* +/− and *Kctd19* KO spermatocytes were immunostained as indicated. Scale bar: 5 μm. The number of MLH1 foci is shown in the scatter plot with median (right). Statistical significance is shown by *p*-value (Mann-Whitney U-test). ****: p < 0.0001. n: number of spermatocytes examined. Asynapsed chromosomes in MLH1+ pachytene spermatocytes are indicated by white arrowheads. **(I)** Chromosome spreads of *Kctd19* +/− and *Kctd19* KO spermatocytes at P21 were immunostained as indicated. Scale bar: 5 μm. Zyg: zygotene, Pac: pachytene, Dip: diplotene. Quantification of meiotic prophase stage per total SYCP3+ spermatocytes of *Kctd19* +/− and *Kctd19* KO mice is shown on the right. n: the number of cells examined. **(J)** Giemza staining of metaphase I chromosomes from *Kctd19* +/− and *Kctd19* KO spermatocytes. **(K)** Immunostaining of squashed metaphase I spermatocytes from *Kctd19* +/− and *Kctd19* KO. The metaphase I kinetochores were identified by MEIKIN immunostaining. Scale bar: 5 μm. Quantification of metaphase I spermatocytes with chromosome misalignment is shown on the right. n: the number of spermatocytes examined. Statistical significance is shown by *p*-value (chi square-test).****: p < 0.0001. **(L)** Seminiferous tubule sections (8-weeks old) were stained for SYCP3, H3S10P and DAPI. Scale bar: 25 μm. M I: Metaphase I spermatocyte. Enlarged images are shown on the rightmost panels. Arrowheads indicate misaligned chromosomes. Shown on the right is the quantification of the seminiferous tubules that have H3S10p+ /SYCP3+ Metaphase I cells per the seminiferous tubules that have SYCP3+ spermatocyte cells in *Kctd19* +/− and *Kctd19* KO testes (bar graph with SD). n: the number of animals examined for each genotype. Statistical significance is shown by *p*-value (t-test). N.S.: Statistically not significant. **(M)** Seminiferous tubule sections from 8-weeks old mice were subjected to TUNEL assay with immunostaining for SYCP3 and H3S10P. Scale bar: 25 μm. Arrowheads indicate TUNEL positive cells. Shown on the right is the quantification of the seminiferous tubules that have TUNEL+ cells per total tubules in *Kctd19* +/− and *Kctd19* KO testes (bar graph with SD). Statistical significance is shown by *p*-values (paired t-test). **: *p* <0.01.

*Kctd19* KO spermatocytes passed through late pachytene as indicated by the presence of H1t staining (Fig. 4G). However, we noticed that the number of MLH1 foci was reduced in *Kctd19* KO pachytene spermatocytes compared to the control (Fig. 4H), suggesting that crossover recombination was partly impaired in the absence of KCTD19. Furthermore, ~50% of *Kctd19* KO MLH1 positive pachytene spermatocytes were accompanied by asynapsed chromosomes (Fig. 4H), suggesting that homolog synapsis was partly defective in *Kctd19* KO. Consistently, diplotene spermatocyte population was reduced compared to WT at P21, the time when the first wave of spermatogenesis reached to diplotene and metaphase I in WT testis (Fig. 4I), suggesting that the progression of meiotic prophase was compromised in *Kctd19* KO. Notably, although *Kctd19* KO spermatocytes reached to metaphase I in adulthood (Fig. 4L), some, if not all, of metaphase I spermatocytes showed chromosome misalignment on the metaphase plate albeit with normal number of bivalent chromosomes with chiasmata (Fig. 4J, K). Therefore, the meiotic prophase defects in *Kctd19* KO derived at least in part from chromosome misalignment at metaphase I in adult male. Furthermore, a subpopulation (31.9%) of the metaphase I cells was TUNEL positive in the *Kctd19* KO seminiferous tubules (Fig. 4M), suggesting that metaphase I cells were eliminated at least in part by apoptosis. Altogether, these results suggested that KCTD19 was required for the completion of meiosis I.

It is worth noting that *Kctd19* KO and *Zfp541* KO mice did not necessarily exhibit the same phenotypes, even though ZFP541 and KCTD19 formed a complex. Whereas spermatocytes beyond pachytene were absent in *Zfp541* KO (Fig. 2), those that reached metaphase I appeared in *Kctd19* KO testis (Fig. 4J, K). Thus, *Zfp541* KO spermatocytes showed severer phenotype than *Kctd19* KO in terms of meiotic prophase progression. Given that ZFP541 remained in the nuclei in *Kctd19* KO whereas both ZFP541and KCTD19 were absent in the nuclei of *Zfp541* KO spermatocytes, ZFP541 may have KCTD19-dependent and independent functions.

### Transcriptome analyses of *Zfp541* KO spermatocytes

Transcriptome analysis on the spermatocytes was performed to address whether the observed phenotype in *Zfp541* KO testes was accompanied by alteration in gene expression. Although the cellular composition of testes changes during development, we assumed the first wave of spermatocyte should progress with same cellular composition in WT and *Zfp541* KO juvenile testes until the defects appear in the mutants. Since ZFP541 appeared from pachytene onward, we analyzed the transcriptomes of meiotic prophase-enriched population that represents the stage when the defects emerge in the mutant testes. For this purpose, we generated *Rec8*-*3xFLAG-HA-p2A-GFP* knock-in (*Rec8*-*3FH-GFP* KI) mice (Fig. S5), as REC8 is expressed exclusively during meiotic prophase (Ishiguro et al., 2011). The meiotic prophase-enriched population was isolated from WT and *Zfp541* KO testes at P18 in *Rec8*-*3FH-GFP* KI background by fluorescent sorting of GFP positive cells (Fig. S5E). We analyzed the transcriptomes of GFP expressing meiotic prophase cells in the control *Zfp541* +/− and *Zfp541* KO by RNA-seq (Fig. 5).

**Figure 5.**
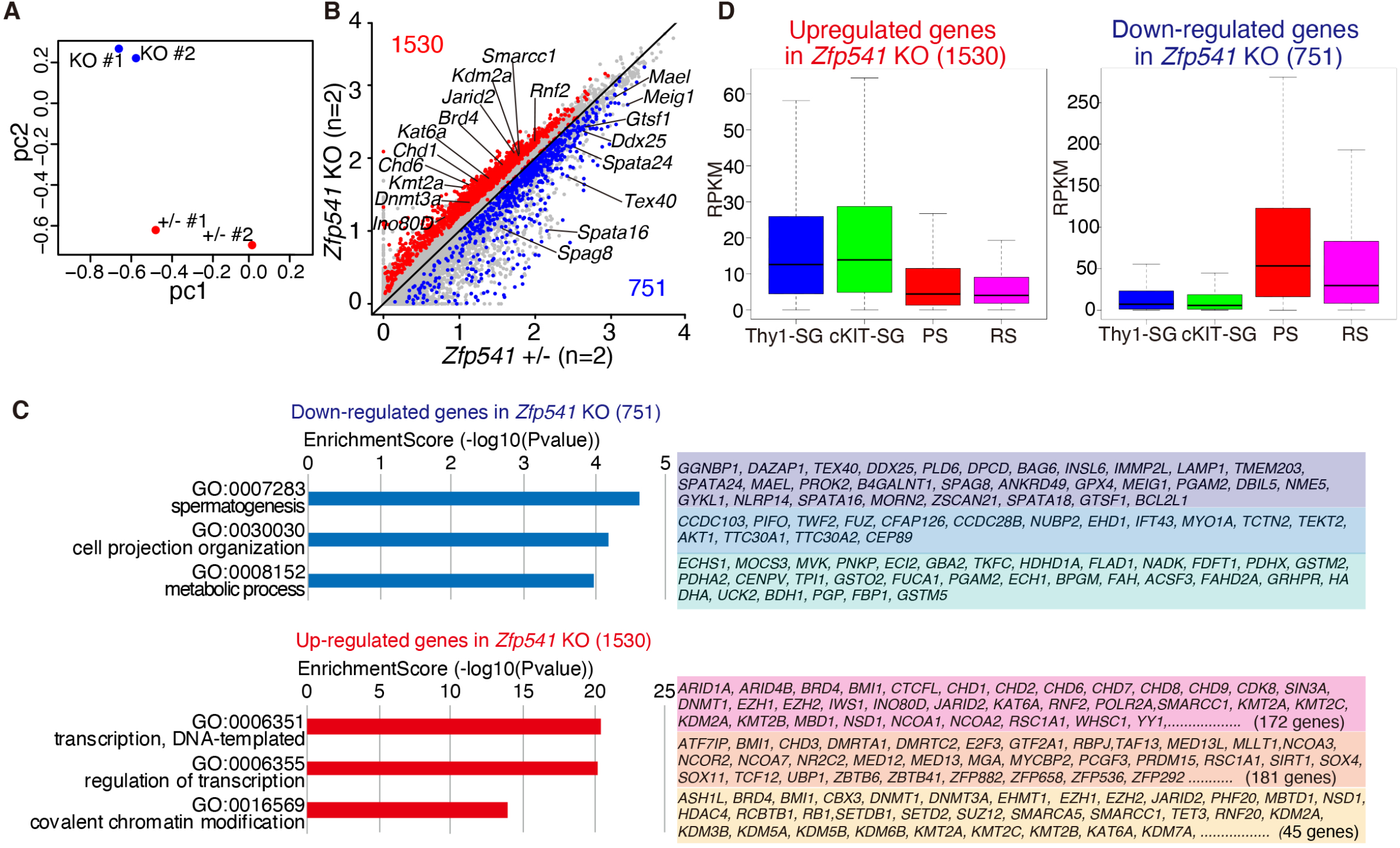
Transcriptome analysis of meiotic prophase spermatocytes in *Zfp541* KO. **(A)** The GFP positive cells were isolated from control *Zfp541*+/− and *Zfp541* KO testes (P18) on the *Rec8*-*3FH-p2A-GFP* KI background by fluorescent sorting. Principal component analysis of the transcriptomes of GFP expressing cells (meiotic prophase spermatocytes) in *Zfp541*+/− and *Zfp541* KO on the *Stra8*-*3FH-p2A-GFP* KI background is shown. **(B)** Scatter plot of the transcriptome of GFP expressing cells in *Zfp541*+/− (*n* = 2) versus *Zfp541* KO (*n* = 2) is shown. The numbers of differentially expressed genes are shown. Significance criteria: false discovery rate ≤ 0.05. **(C)** GO analysis of the downregulated genes (upper) and upregulated genes (lower) in GFP expressing cells of *Zfp541* KO testes. See Table S1 for complete gene list of the GO analyses. **(D)** Expression levels (RPKM) of the upregulated 1530 genes (left) and the downregulated 751 genes (right) in GFP expressing spermatocytes of *Zfp541* KO are shown by box plot with average. The upregulated and downregulated genes in *Zfp541* KO were reanalyzed with the previously published data of stage-specific expression (Hasegawa et al., 2015; Maezawa et al., 2018b). Thy-SG: Thy+ spermatogonia, cKIT-SG: cKIT+ spermatogonia, PS: pachytene spermatocytes, RS: round spermatid.

Principal component analysis (PCA) revealed that the overall transcriptomes of the GFP positive cells in *Zfp541* KO testes were different from those of the control testes at P18 (Fig. 5A, B). Notably, gene ontology (GO) analysis of differentially expressed genes (DEG) between P18 control *Zfp541* +/− and *Zfp541* KO testes indicated that the genes involved in transcriptional regulation and covalent chromatin modification were upregulated in *Zfp541* KO (Fig. 5C, Table S1). Reanalysis of the data together with the data from previous studies on transcriptomes during spermatogenesis (Hasegawa et al., 2015; Maezawa et al., 2018b) demonstrated that the upregulated genes in *Zfp541* KO (1530 genes) were overall expressed in spermatogonia and the expression declined from prophase onward (Fig. 5D). In contrast, the downregulated genes in *Zfp541* KO (751 genes), were overall less expressed in spermatogonia and highly expressed in prophase spermatocytes (Fig. 5D). Notably, GO analysis demonstrated that the genes involved in spermatogenesis were downregulated in *Zfp541* KO (Fig. 5C, Table S1), which at least in part accounts for the aforementioned cytological observation showing that *Zfp541* KO spermatocytes failed to exit meiotic prophase. Thus, these results suggest that ZFP541 is required for the completion of the meiotic prophase program in male.

### ZFP541 binds to the promoter regions of the genes associated with transcriptional regulation

ZFP541 is assumed to possess putative DNA-binding domains and associate with histone deacetylases, implying it plays a role in the regulation of transcription via the modulation of chromatin status. So we further investigated the ZFP541-target sites on the genome by chromatin immunoprecipitation followed by sequencing (ChIP-seq) analysis. ZFP541 binding sites were defined by commonly identified sites in two independent ChIP-seq data sets using two different antibodies (Fig S6A-C). As a result, ZFP541 bound to 6,135 sites (5,921 nearest genes, which were assigned regardless of the distance from the ZFP541-binding sites), of which 32.3 % and 26.1% resided around the gene promoter regions and the 5’UTR on the mouse genome, respectively (Fig. 6A). Overall, ZFP541-binding sites resided within +/− 2 kb of the TSS (4,689 genes) (Fig. 6B, Fig. S6B, C, Table S1). Notably, DNA binding motif analysis indicated that ZFP541-ChIP enriched GC rich DNA sequences (Fig. 6C), suggesting that ZFP541 directly binds to the promoters through these motifs. Furthermore, the ZFP541-bound sites largely showed increased repressive histone mark H3K27me3 and reciprocally decreased active histone H3K27ac during the transition from prophase to round spermatids (Maezawa et al., 2018b) (Maezawa et al., 2020) (Fig. S6D), suggesting that ZFP541-bound sites overall undergo repressive states after the exit from meiotic prophase. It should be mentioned that ZFP541-bound sites were less presented in the X and Y chromosomes, consistent with the immunostaining of ZFP541 that showed absence of staining in the XY body (Fig. S6E, F).

**Figure 6.**
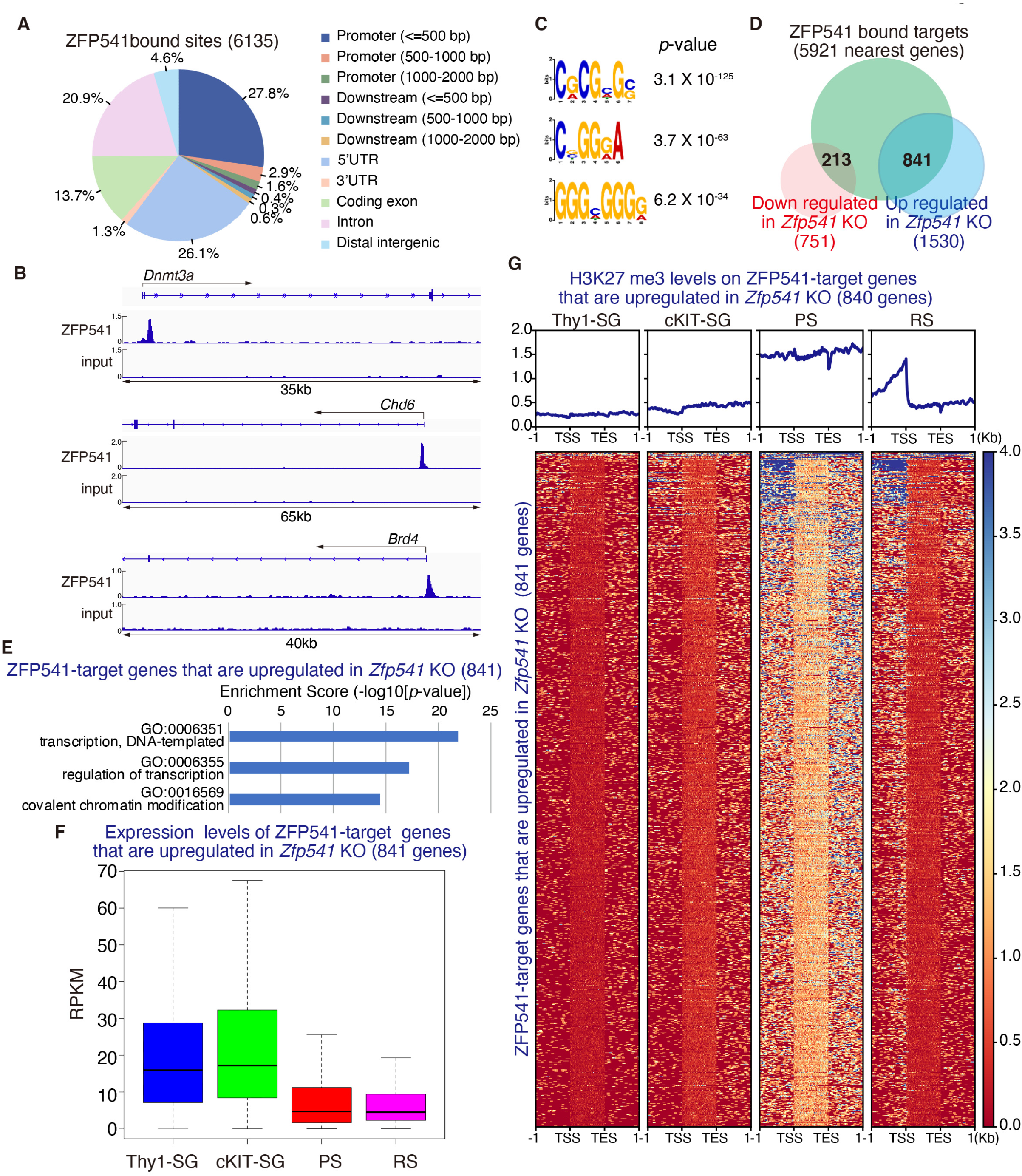
ZFP541 binds to the gene promoter regions. **(A)** ZFP541 binding sites were classified by the genomic locations as indicated. **(B)** Genomic view of ZFP541 ChIP-seq and input DNA data over representative gene loci. Genomic coordinates were obtained from RefSeq. **(C)** Top 3 sequence motifs enriched in ZFP541 ChIP-seq with *p*-values. **(D)** Venn diagram representing the overlap of ZFP541-bound genes (5923 nearest genes), downregulated (751 genes), and upregulated genes (1530 genes) in *Zfp541* KO mice. **(E)** GO analysis of the ZFP541-bound genes that were upregulated in *Zfp541* KO mice (840 genes) (p < 1.0 × 10^−6^). Top 3 biological processes ranked by log (*p*-value) are listed. The ZFP541-bound genes that were downregulated in *Zfp541* KO (213genes) showed no GO terms statistically enriched. See Table S2 for complete gene list of the GO analyses. **(F)** Expression levels (RPKM) of the ZFP541-bound genes that were upregulated in *Zfp541* KO spermatocytes (840 genes) are shown by box plot with average, as in Figure 5D. Thy-SG: Thy+ spermatogonia, cKIT-SG: cKIT+ spermatogonia, PS: pachytene spermatocytes, RS: round spermatid. **(G)** Heat map of H3K27me3 levels on the ZFP541-target genes that are upregulated in *Zfp541* KO (840 genes). H3K27me3 levels are shown on the genomic regions between −1.0 kb upstream of TSS and +1.0□kb downstream of TES. Color key is shown. Average distributions of H3K27me3 are shown (upper).

Among the ZFP541-bound targets, 841 and 213 genes were identified within the upregulated genes and the down regulated genes in *Zfp541* KO GFP positive cells, respectively (Fig. 6D), supporting the idea that ZFP541 regulates the expression of those target genes. Remarkably, GO analysis revealed that the ZFP541-target genes upregulated in *Zfp541* KO (841 genes) such as *Dnmt1, Dnmt3A* (DNA methyl transferase), *Ino80D, Chd1, Chd2, Chd3, Chd6* (chromatin remodeling), *Kat6A, Kmt2C, Kmt2B, Kdm2A, Kdm5B, Ash1L, Bmi1, Jarid2, Ezh1, Ezh2, Ehmt1, Rnf2, Suz12* (Histone modification), and *Ctcfl* (chromatin binding) were associated with the biological processes of transcriptional regulation and covalent chromatin modification (Fig. 6E, Table S2). In contrast, among the ZFP541-target genes downregulated by *Zfp541* KO (213 genes), we did not find GO terms that showed statistical significance (Table S2).

Since ZFP541 was associated with histone deacetylases, we analyzed the ZFP541-bound genes along with the RNA expression and histone modification levels during spermatogenesis using the data set from previous studies (Hasegawa et al., 2015; Maezawa et al., 2018b). Strikingly, overall expression levels of those ZFP541-target genes that were upregulated in *Zfp541* KO (841 genes) were declined in prophase spermatocytes and round spermatids (Fig. 6F). Accordingly, overall H3K27me3 levels were increased across the gene body in prophase spermatocytes and remained high at the promoter regions in round spermatids (Fig. 6G, Fig S6D). This suggests that the HDAC1/2-containing ZFP541 complex directly represses the transcription of a subset of critical genes that are involved in a broad range of transcriptional regulation and chromatin modification prior to the exit from meiotic prophase. These findings suggest that the ZFP541 complex binds to and regulates the expressions of a subset of late meiotic genes for pachytene exit (Fig. 7).

**Figure 7.**
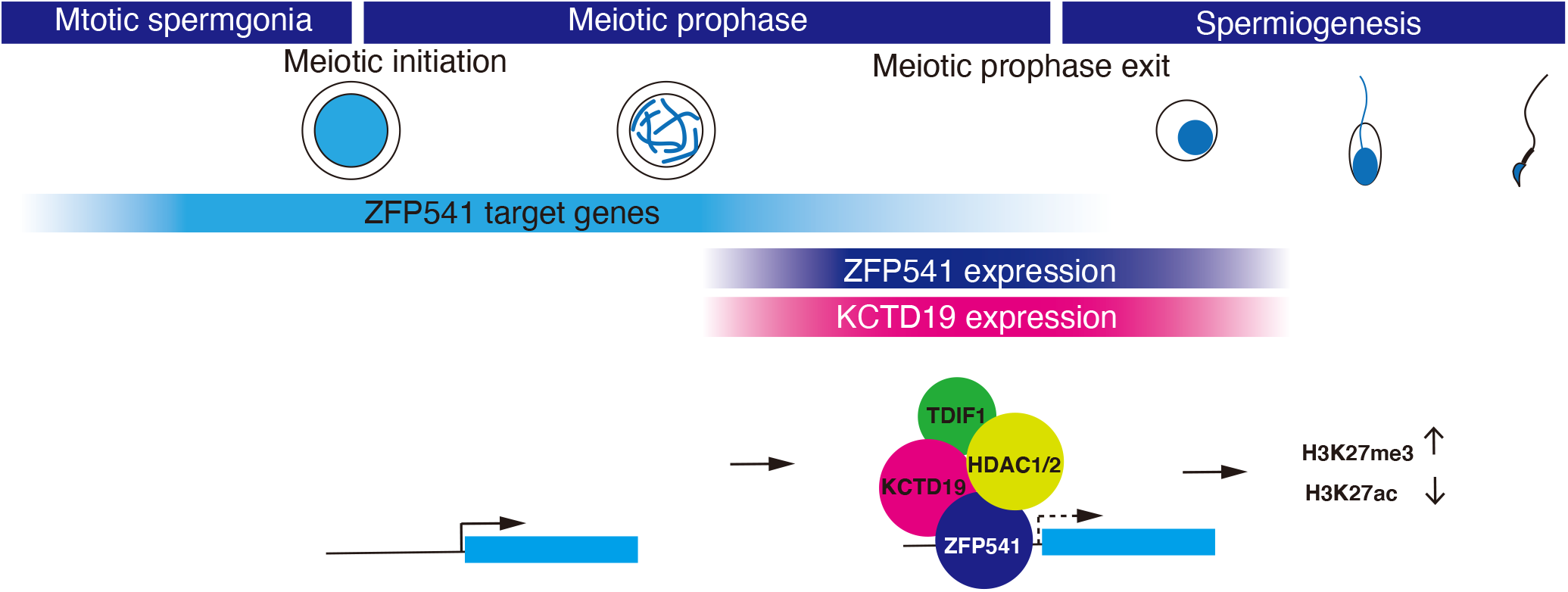
The ZFP541-KCTD19 complex is required for meiotic prophase exit. Schematic model of the ZFP541-KCTD19 complex that suppresses ZFP541-target genes prior to prophase exit.

## Discussion

During spermatogenesis, meiosis is followed by spermiogenesis, which is accompanied by robust alterations of transcriptional program and chromatin status to prepare for sperm production. In the present study, we demonstrate that ZFP541 forms a complex with KCTD19, HDAC and TDIF1 in spermatocytes and round spermatids (Fig.3A, Figs S2–S3). Our genetic study demonstrated that both ZFP541 and KCTD19, whose expressions are germ cell specific, are required for the completion of meiotic prophase I in spermatocytes (Figs. 2, 4), highlighting the sexual difference in the transcriptional program of meiotic prophase. Interestingly, although ZFP541 and KCTD19 forms a complex, *Zfp541* KO and *Kctd19* KO mice exhibited different phenotypes (Figs. 2, 4). Whereas spermatocyte beyond pachytene was absent in *Zfp541* KO (Fig. 2), metaphase I cells appeared in *Kctd19* KO testis (Fig. 4). Thus, *Zfp541* KO spermatocytes showed severer phenotype in meiotic prophase progression than *Kctd19* KO. Because both ZFP541and KCTD19 were absent in the nuclei of *Zfp541* KO spermatocytes, whereas ZFP541 remained in the nuclei and was functional in *Kctd19* KO (Fig. 4C), ZFP541 may have a function independent of KCTD19.

It should be mentioned that MIDEAS and Transcription regulatory factor 1 (TRERF1) as well as TDIF1 were identified in KCTD19 imuunoprecipitates (Fig. S3B). Previously, it was shown that TDIF1, MIDEAS and HDAC1 formed a mitotic deacetylase (MiDAC) complex in mitotic cells, and that loss of either TDIF1 or MIDEAS led to chromosome misalignment in mitosis (Turnbull et al., 2020). Furthermore, MIDEAS and TRERF1 share homology in their ELM/SANT domains, and MIDEAS and TRERF1 interact with each other (Hein et al., 2015) (Turnbull et al., 2020). Given that MIDEAS and TRERF1, as well as ZFP541, possess ELM/SANT domains, KCTD19 may form a subcomplex that consists of MIDEAS, TRERF1, TDIF1 and HDAC1/2, which is distinct from the ZFP541-KCTD19-TDIF1-HDAC1/2 complex. Therefore, it is possible that the defect in metaphase I chromosome alignment in *Kctd19* KO was indirectly caused by the lack of MIDEAS-TRERF1-KCTD19-TDIF1-HDAC1/2 subcomplex, rather than ZFP541-KCTD19-TDIF1-HDAC1/2 complex (Fig. 4K).

Crucially, our ChIP-seq analysis indicates that ZFP541 binds to a broad range of genes associated with biological processes of transcriptional regulation and covalent chromatin modification, such as DNA methylation, chromatin remodeling, and histone modification (Figs. 6B, E). Because ZFP541 associates with HDAC1 and HDAC2 (Fig.3A, Fig. S2), we reasoned that ZFP541 plays a role in repressing the target genes prior to the completion of meiotic prophase program (Fig. 7). Consistent with this idea, the expression levels of the ZFP541-target genes were repressed and were accompanied by repressive H3K27me3 mark in round spermatids (Figs. 6F, G). Furthermore, it should be mentioned that most of the ZFP541-target genes, if not all, were well overlapped with the target genes of a germ cell specific PRC1 component, SCML2 in germline stem cells (Fig. S6G) (Hasegawa et al., 2015) (Maezawa et al., 2018a) (Maezawa et al., 2018b) Since SCML2 binds somatic/progenitor genes in the stem cell stage to suppress these genes after meiosis (Hasegawa et al., 2015), it is possible that those ZFP541-target genes might be pre-marked with SCML2 and Polycomb proteins for later suppression of transcription. In support of this possibility, ZFP541-target genes are marked with PRC2-mediated H3K27me3 (Fig. 6G), which is established in downstream of SCML2 in pachytene spermatocytes (Maezawa et al., 2018b). Therefore, we concluded that ZFP541 triggers the reconstruction of the transcription network to promote the exit from prophase, finalize meiotic divisions, and proceed into spermatid production, which may be similar to the function of the transcription factor Ndt80 that promotes pachytene exit and spore formation in budding yeast (Xu et al., 1995) (Chu and Herskowitz, 1998). Our study should shed light on the regulatory mechanisms of gene expression that promotes meiotic prophase pachytene exit leading to spermatid differentiation.

## Acknowledgments

The authors thank Drs. Masahito Ikawa, Seiya Oura (Osaka university) for discussion and sharing unpublished data, Sally Fujiyama (The university of Tokyo), Etsushi Kitamura, Kaho Okamura (Kumamoto University) for technical support, and Marry Ann Handel for provision of H1t antibody. This work was supported in part by KAKENHI (#19K06642); Takeda Science Foundation (to Y.T.), and KAKENHI (#17H03634, #18K19304, #19H05245, #19H05743, #20H03265, #20K21504, #JP 16H06276) from MEXT, Japan; Joint Usage and Joint Research Programs from the Institute of Advanced Medical Sciences, Tokushima University (30-A-7); Grants from The Sumitomo Foundation, The Naito Foundation, Astellas Foundation for Research on Metabolic Disorders, Daiichi Sankyo Foundation of Life Science, The Uehara Memorial Foundation (to K.I.).

## Author contributions

YT CK KT performed the mice cytology experiments. KT KI NT performed MS analyses. YT performed the ChIP-seq, supported by KM HN RM MT. YT performed the RNA-seq, supported by SU. KH AS KM supported data analyses of ChIP-seq and RNA-seq. AS SN performed reanalysis of RNA-seq and ChIP-seq data. YT TA performed the ATAC-seq. SF performed histological analyses. KA KI designed the knockout mice. KI supervised, conducted the study and wrote the manuscript.

## Figure legend

**Supplementary Figure 1.**
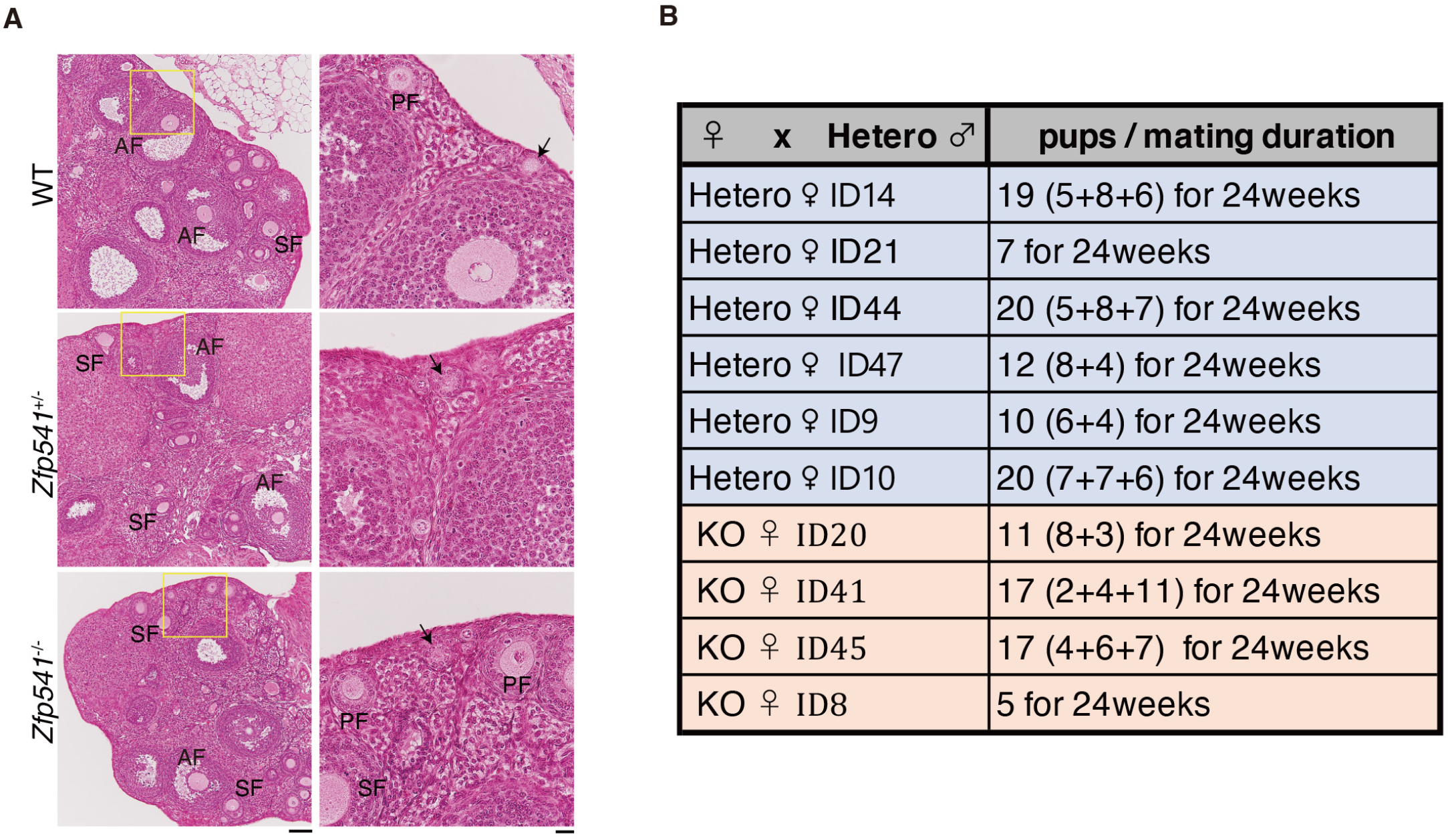
Phenotypic analyses of *Zfp541* KO ovaries (related to Figure 2) **(A)** Hematoxylin and eosin staining of the sections from WT, *Zfp541* +/− and *Zfp541* KO ovaries (8-weeks old). Scale bar: 100 μm. Enlarged images are shown on the right. Scale bar: 20 μm. Arrow: primordial follicle. PF: primary follicle, SF: secondary follicle, AF: antral follicle. **(B)** Fertility of *Zfp541* KO females was examined by mating with *Zfp541* +/− males for the indicated period.

**Supplementary Figure 2.**
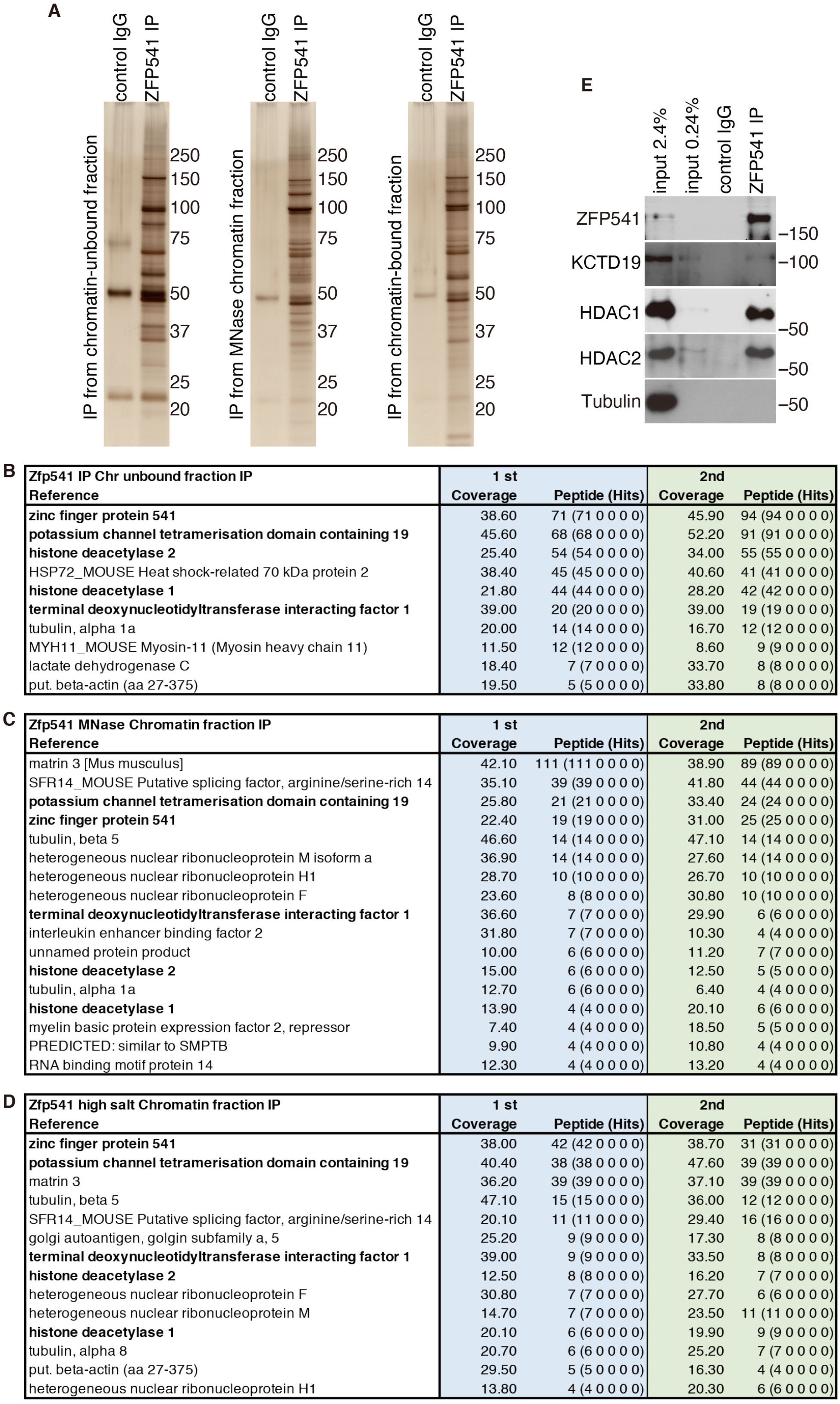
MS analyses of ZFP541 interacting factors in testis extracts (related to Figure 3) **(A)** Silver staining of the immunoprecipitates by anti-ZFP541 antibody (ZFP541-IP) from the chromatin-unbound, MNase released chromatin, and chromatin-bound fractions of the testis extracts. **(B-D)** The immunoprecipitates were subjected to liquid chromatography tandem-mass spectrometry (LC-MS/MS) analyses. The proteins identified by the LC-MS/MS analyses of ZFP541-IP are presented after excluding the proteins detected in the control IgG-IP. The proteins, which were reproducibly identified by two independent LC-MS/MS analyses (1st and 2nd) with more than 3 different peptide hits, are listed with the number of peptide hits and % coverage in the table. Shown are the identified proteins of ZFP541 immunoprecipitates from chromatin-unbound (B), MNase released chromatin (C), and chromatin-bound (D) fractions of the testis. The proteins reproducibly identified in the three different fractions are shown in bold. **(E)** Immunoblot showing the immunoprecipitates of ZFP541 from chromatin extracts of WT mouse testes.

**Supplementary Figure 3.**
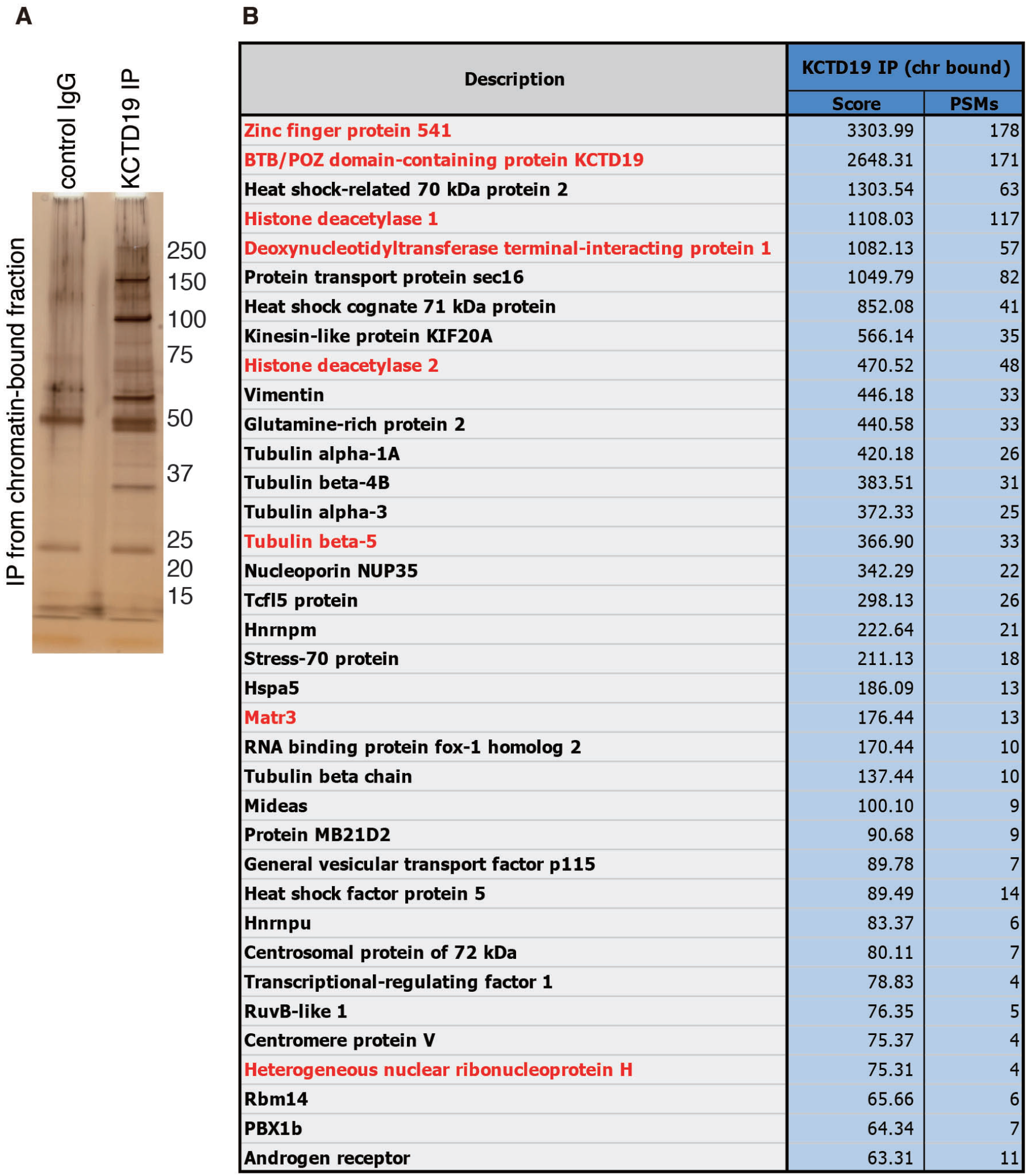
MS analyses of KCTD19 interacting factors in testis extracts (related to Figure 3) **(A)** Silver staining of the immunoprecipitates by anti-KCTD19 antibody (KCTD19-IP) from the chromatin-bound fraction of the testis extracts. **(B)** The immunoprecipitates from the chromatin-bound fraction of the testis extracts were subjected to liquid chromatography tandem-mass spectrometry (LC-MS/MS) analyses. The proteins identified by the LC-MS/MS analysis of KCTD19-IP are presented after excluding the proteins detected in the control IgG-IP. The proteins with more than 3 different peptide hits are listed with the number of peptide hits and Mascot scores. The proteins commonly identified in KCTD19-IP and ZFP541-IP from the chromatin-bound fraction of the testis extracts are shown in red.

**Supplementary Figure 4.**
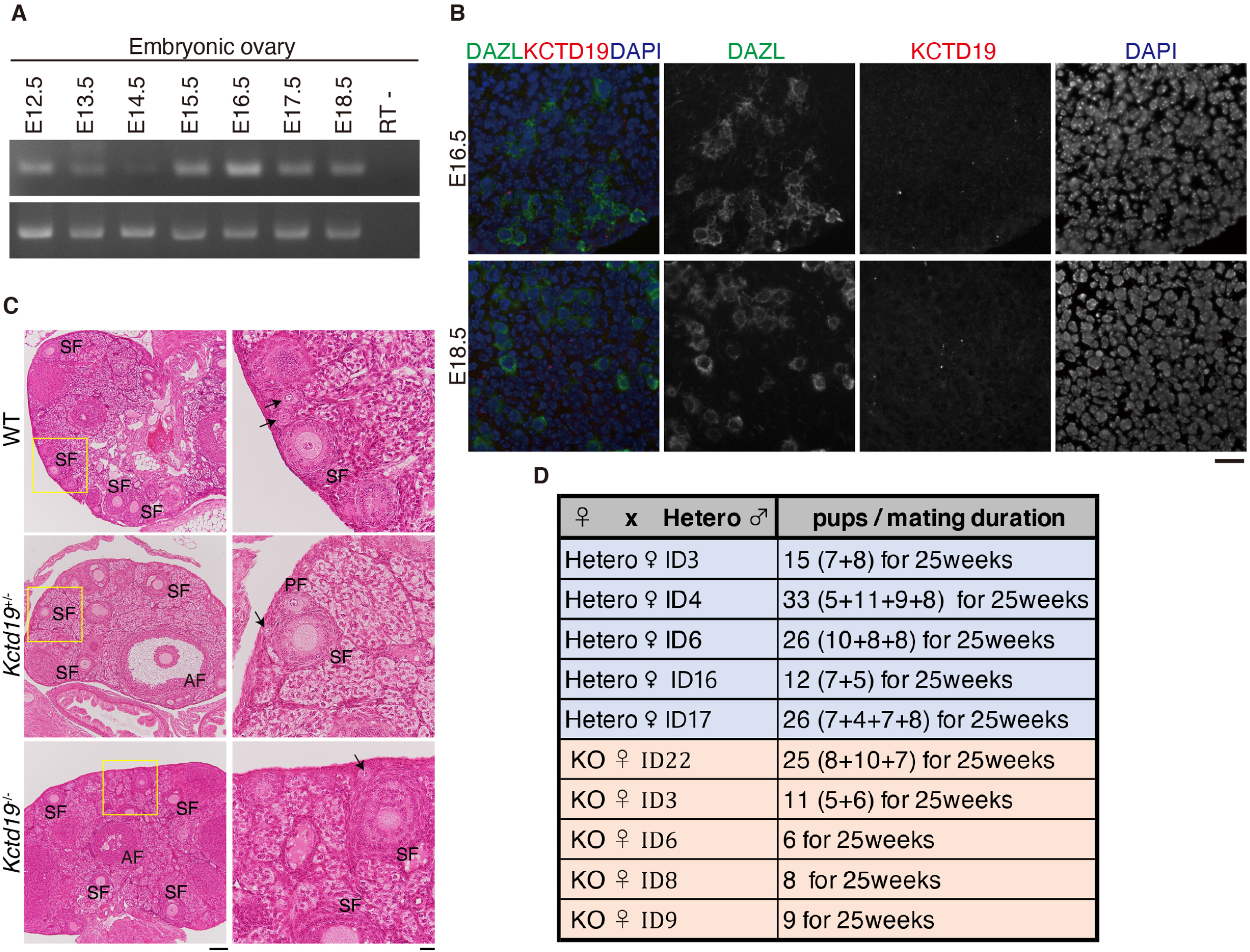
Phenotypic analyses of *Kctd19* KO ovaries (related to Figure 4) **(A)** RT-PCR analysis of *Kctd19* in embryonic ovaries (E12.5-18.5). RT- indicates control PCR without reverse transcription. **(B)** Embryonic ovary sections (E16.5 and E18.5) were stained for KCTD19, DAPIand a germ cell marker DAZL. Scale bars: 25 μm. **(C)** Hematoxylin and eosin staining of the sections from WT, *Kctd19* +/− and *Kctd19* KO ovaries (8-weeks old). Scale bar: 100 μm. Enlarged images are shown on the right. Scale bar: 20 μm. Arrow: primordial follicle, PF: primary follicle, SF: secondary follicle, AF: antral follicle. **(D)** Fertility of *Kctd19* KO females was examined by mating with *Kctd19* +/− males for the indicated period.

**Supplementary Figure 5.**
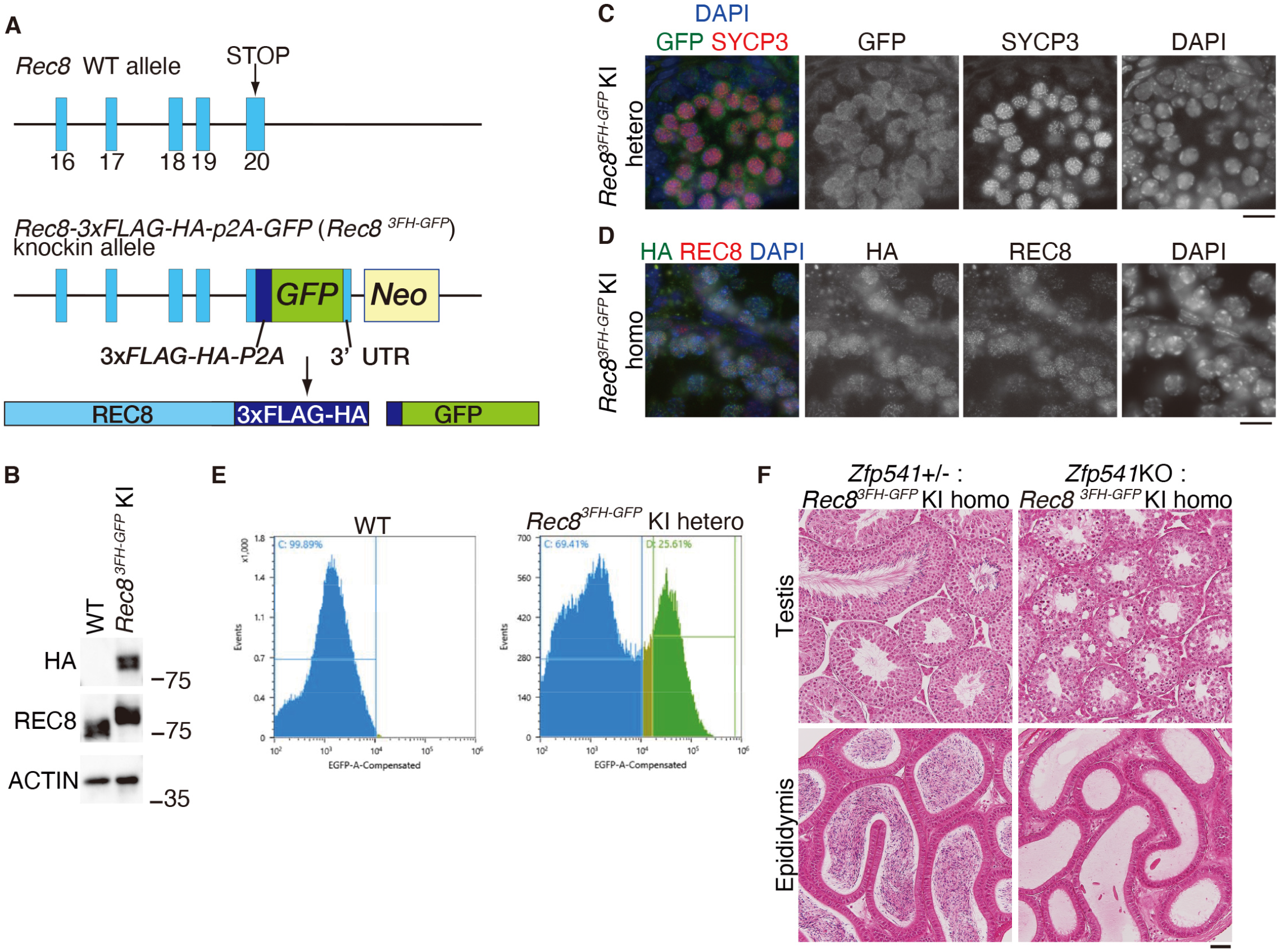
Generation of *Rec8-3xFLAG-HA-p2A-GFP* knock-in mice (related to Figure 5) **(A)** Schematic illustrations of the *Rec8* WT allele and the *Rec8-3xFLAG-HA-p2A-GFP* knock-in (*Rec8-3FH-GFP* KI) allele. Blue boxes represent exons. Coding exon 20 is followed by *3xFLAG-HA-p2A-GFP* and the 3’UTR. Neo: Neomycine resistance gene. **(B)** Immunoblot of testis extracts from WT (non-tagged control) and the *Rec8-3xFLAG-HA-p2A-GFP* KI homozygous mice. Note that anti-REC8 blot detected 3xFLAG-HA tagged REC8 at higher molecular weight than the endogenous REC8. **(C)** Seminiferous tubule sections from *Rec8*-*3xFLAG-HA-p2A-GFP* KI heterozygous testis were stained for SYCP3, GFP and DAPI. Scale bar: 15 μm. **(D)** Seminiferous tubule sections from *Rec8*-*3xFLAG-HA-p2A-GFP* KI homozygous testis were stained for HA, REC8 and DAPI. Scale bar: 15 μm. **(E)** The meiotic prophase spermatocytes were isolated by fluorescent sorting of GFP positive cells from testes on the *Stra8*-*3FH-p2A-GFP* KI background. **(F)** Hematoxylin and eosin staining of the testes (upper) and epididymis (lower) sections from *Zfp541+/−* and *Zfp541* KO in the *Rec8-3HA-GFP* KI homozygous background (8-weeks old). Scale bar: 50 μm. Note that the REC8-3xFLAG-HA fusion protein localizes along the axes in a similar manner to that observed in normal WT testis. The fusion protein was physiologically functional considering that homozygous male and female mice with the KI allele showed normal fertility.

**Supplementary Figure 6.**
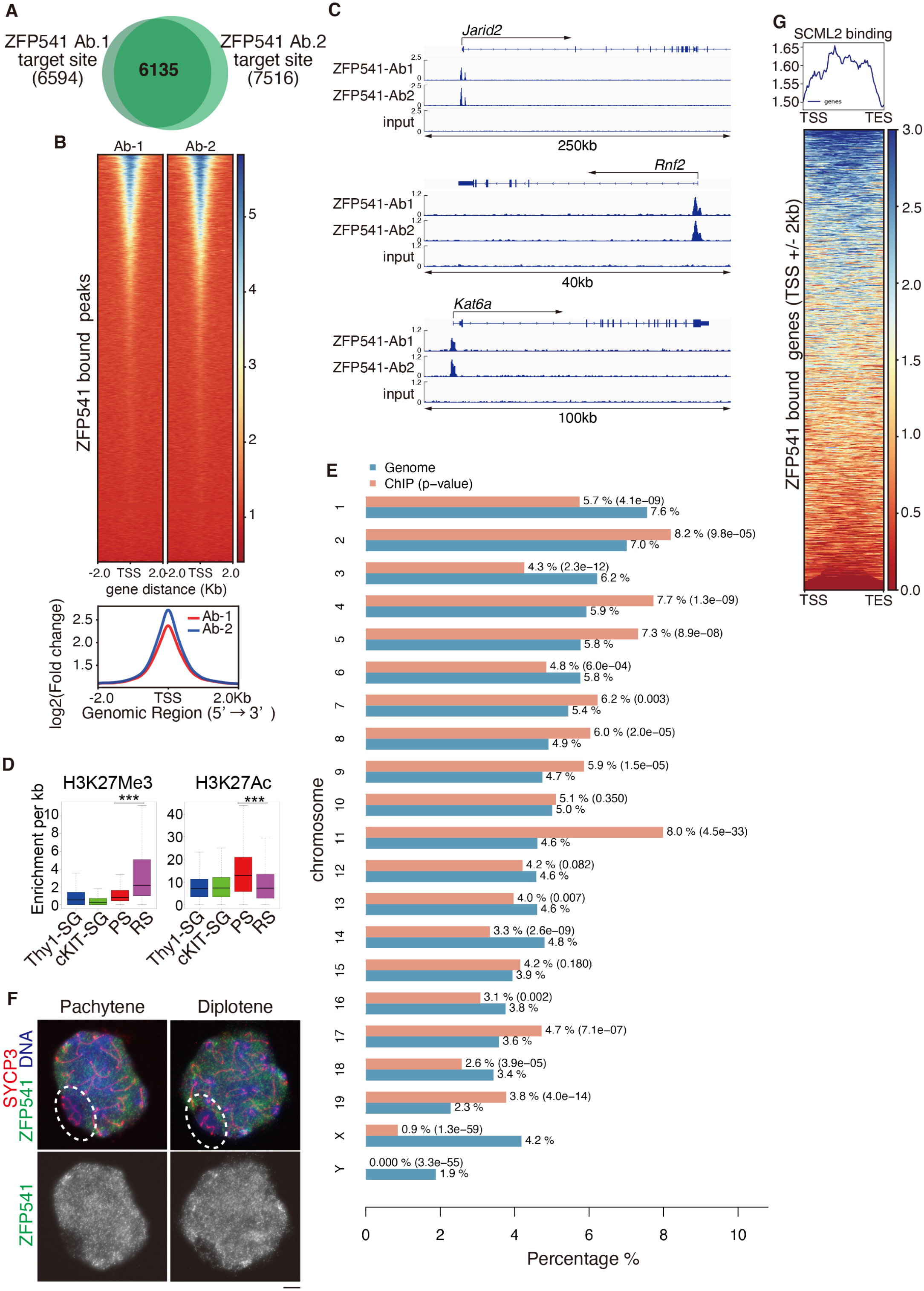
ChIP-seq analysis of ZFP541 in the testis (related to Figure 6) **(A)** Venn diagram representing the overlap of ZFP541-bound sites (6135 sites) from two independent ZFP541 ChIP-seq data sets using two different antibodies. **(B)** Heat map of the common ZFP541 binding sites (6135 sites) of two independent ZFP541 ChIP-seq at the positions −2.0 kb upstream to +2.0◻kb downstream relative to the TSS. Average distributions of ZFP541-binding peak for two independent ZFP541 ChIP-seq are shown on the bottom. **(C)** Genomic view of ZFP541 ChIP-seq (duplicates using two different anti-ZFP541antibodies, Ab-1 and Ab-2) and input DNA data over representative gene loci. Genomic coordinates were obtained from RefSeq. **(D)** Enrichment of H3K27me3 (left) and H3K27Ac (right) levels on the whole ZFP541-bound sites (6135 sites/5923 nearest genes) during spermatogenesis are shown by box plot with average. Thy-SG: Thy+ spermatogonia, cKIT-SG: cKIT+ spermatogonia, PS: pachytene spermatocytes, RS: round spermatid. *** p < 2.2 × 10 ^−16^ (Wilcoxon rank sum test). ChIP-seq data of H3K27me3 and H3K27Ac are derived from previous studies (Maezawa et al., 2018b) (Maezawa et al., 2020). **(E)** Distribution of ZFP541 ChIP-seq peaks on the chromosomes. Proportion (%) of the chromosome which were assigned with ZFP541 ChIP-seq peaks are shown with *p*-values. Note that ZFP541 ChIP-seq peaks were excluded from the Y chromosome, and were less presented in the X chromosome. **(F)** Pachytene and diplotene spermatocyte nuclei were immunostained for SYCP3 and ZFP541. The XY body is encircled by dashed lines. Scale bar: 5 μm. **(G)** Heat map of SCML2 levels (Hasegawa et al., 2015) on the ZFP541-target genes that are within +/− 2kb of TSS (4,689 genes). SCML2 levels are shown on the genomic regions between TSS and TES. Color key is shown. Average distributions of SCML2 are shown (upper).

## STAR Methods

### Lead Contact and Materials Availability

#### Data and materials availability

Further information and requests for resources and reagents should be directed to and will be fulfilled by the Lead Contact, Kei-ichiro Ishiguro. *Zfp541* and *Kctd19* knockout mouse and *Rec8*-*3xFLAG-HA-p2A-GFP* knock-in mouse lines generated in this study have been deposited to Center for Animal Resources and Development (CARD, ID 2857 for *Zfp541*, ID 2879, ID 2880 for *Kctd19* Line #12, Line #26 respectively, ID2681 for *Rec8*-*3xFLAG-HA-p2A-GFP*). The ZFP541 ChIP-seq data of mouse testes are deposited in the GEO Sequence Read Archive (SRA) under accession number GSE163916. RNA-seq data is deposited under GSE163917

. The antibodies are available upon request. There are restrictions to the availability of antibodies due to the lack of an external centralized repository for its distribution and our need to maintain the stock. We are glad to share antibodies with reasonable compensation by requestor for its processing and shipping. All unique/stable reagents generated in this study are available from the Lead Contact with a completed Materials Transfer Agreement.

### Experimental Model and Subject Details

#### Animal experiments

*Zfp541* and *Kctd19* knockout mice were C57BL/6 background.

*Rec8*-*3xFLAG-HA-p2A-GFP* knock-in mice were congenic with the C57BL/6 background. Whenever possible, each knockout animal was compared to littermates or age-matched non-littermates from the same colony, unless otherwise described. Animal experiments were approved by the Institutional Animal Ethics Committees of Kumamoto University (approval F28-078, A2020-006, A30-001, A28-026).

#### Generation of *Zfp541* knockout mice and genotyping

*Zfp541* knockout mice were generated by introducing Cas9 protein (317-08441; NIPPON GENE, Toyama, Japan), tracrRNA (GE-002; FASMAC, Kanagawa, Japan), synthetic crRNA (FASMAC), and ssODN into C57BL/6N fertilized eggs using electroporation. For the generation of *Zfp541* Exon1-13 deletion (Ex1-13Δ) allele, the synthetic crRNAs were designed to direct CATGGAGCCATACAGTCTT(GGG) of the *Zfp541* exon 1 and ACGACCAAAGAAACAGTGCT(GGG) in the intron 13. ssODN: 5’-GAACGAATTCACCTTCTAGCCAAAAACGACCAAAGAAACAGTatttatttACTGTATGGCTCCATGGCTTTCCACTGCCAGGTCCTGGCT-3’ was used as a homologous recombination template.

The electroporation solutions contained [10μM of tracrRNA, 10μM of synthetic crRNA, 0.1 μg/μl of Cas9 protein, ssODN (1μg/μl)] for *Zfp541* knockout in Opti-MEM I Reduced Serum Medium (31985062; Thermo Fisher Scientific). Electroporation was carried out using the Super Electroporator NEPA 21 (NEPA GENE, Chiba, Japan) on Glass Microslides with round wire electrodes, 1.0 mm gap (45-0104; BTX, Holliston, MA). Four steps of square pulses were applied (1, three times of 3 mS poring pulses with 97 mS intervals at 30 V; 2, three times of 3 mS polarity-changed poring pulses with 97 mS intervals at 30 V; 3, five times of 50 mS transfer pulses with 50 mS intervals at 4 V with 40% decay of voltage per each pulse; 4, five times of 50 mS polarity-changed transfer pulses with 50 mS intervals at 4 V with 40% decay of voltage per each pulse).

The targeted *Zfp541* Ex1-13Δ allele in F0 mice were identified by PCR using the following primers; Zfp541-F1: 5’-AGCTAGCTGCCAGCGAGGGCTCTTC-3’ and Zfp541-R3: 5’-GAGGCAGCAGAAGGGAGGTAGGATG-3’ for the knockout allele (404bp). Zfp541-F1 and Zfp541-R2: 5’-GGTTGAGTGTGTCACTGCAGTTGAG-3’ for the Ex1 of WT allele (199 bp). Zfp541-F2: 5’-AACGTAGGAAGCAGATTTCAGGCGG −3’ and Zfp541-R1: 5’-TCCCTCAGCTGGGCCATCCAAGTCC −3’ for the Ex10-11 of WT allele (613 bp). The PCR amplicons were verified by Sanger sequencing. Primer sequences are listed in Table S3.

#### Generation of *Kctd19* knockout mice and genotyping

*Kctd19* knockout mouse was generated as described above. For the generation of *Kctd19* Exon3-12 deletion (Ex3-12Δ) allele, the synthetic crRNAs were designed to direct GAAGTGTATCAGCAAGCCCT (CGG) of the *Kctd19* exon 3 and AAGAGGCCAATCATCTAGGG (TGG) in the intron 12. ssODN: 5’-ACCCTAGATGATTGGCCTCTTGGTGGCACCTCTGCCTCCC tttatttattcaGCTTGCTGATACACTTCCATGTCCGCCAGTAGTTCAGGGATGCT −3’ was used as a homologous recombination template.

The targeted *Kctd19* Ex3-12Δ allele in F0 mice were identified by PCR using the following primers; Kctd19-F1: 5’-AGGCAGCACTCTTCTCTGTGGGACAG-3’ and Kctd19-R1: 5’-TTTCCCATGCCACCTGTGCCTTTCC-3’ for the knockout allele (279bp). Kctd19-F1 and Kctd19-R2: 5’-TGCCCGTGACGGTACCTGTGAAGGC-3’ for the Ex3 of WT allele (192 bp). Kctd19-F2: 5’-GGCAAGCAGAGGCCCAAGGACAGAG −3’ and Kctd19-R1 for the intron12 of WT allele (714 bp). The PCR amplicons were verified by Sanger sequencing. Primer sequences are listed in Table S3.

#### Generation of *Rec8*-*3xFLAG-HA-p2A-GFP* knock-in mouse and genotyping

The targeting vector was designed to insert 3xFLAG-HA-p2A-EGFP-3’UTR in frame with the coding sequence into the Exon 20 of the *Rec8* genomic locus. Targeting arms of 1018bp and 812bp fragments, 5’ and 3’ of the Exon 20 of *Rec8* gene respectively, were generated by PCR from mouse C57BL/6 genomic DNA and directionally cloned flanking p*GK-Neo*-polyA and *DT-A* cassettes. The 5’ arm is followed by nucleotide sequences encoding 3xFLAG, HA, p2A, EGFP and the 3’UTR of *Rec8* gene. TT2 ES (Yagi et al., 1993) cells were co-transfected with the targeting vector and pX330 plasmids (Addgene) expressing Crispr-gRNAs directing TATGTGTTCAGAGCTGAGAC(tgg) and GGAAGGGAGGGGCTGCGCTG(agg), which locates at the 3’ region of the Exon 20 of *Rec8* gene. The G418-resistant ES clones were screened for homologous recombination with the *Rec8* locus by PCR using primers Rec8_5arm_F1: 5’-GGAGCTCTTCAGAACCCCAACTCTC-3’ and KI96ES-19814R-HA: 5’-GGGCACGTCGTAGGGGTATCCCTTG −3’ for the left arm (1490 bp).

The homologous recombinant cells were isolated and chimeric mice were generated by aggregation (host ICR) of recombinant ES cells. Chimeric males were mated to C57BL/6N females and the progenies were genotyped by PCR using the primers Rec8-Left2-F: 5’-TGAGCAAGTTCATTAAACCATATCCTG −3’ and Rec8-Rightarm-R: 5’-CTCTTGAAGCTGACATCTTTGGTTAC −3’ for the knock-in allele (2844 bp) and the WT allele (1045bp). Primer sequences are listed in Table S3.

#### PCR with reverse transcription

Total RNA was isolated from tissues and embryonic gonads using TRIzol (Thermo Fisher). cDNA was generated from total RNA using Superscript III (Thermo Fisher) followed by PCR amplification using Ex-Taq polymerase (Takara) and template cDNA. For RT-qPCR, total RNA was isolated from WT and *Meiosin* KO testes, and cDNA was generated as described previously (Ishiguro et al., 2020). *Zfp541* cDNA was quantified by ΔCT method using TB Green Premix Ex Taq II (Tli RNaseH Plus) and Thermal cycler Dice (Takara), and normalized by *GAPDH* expression level.

Sequences of primers for RT-PCR are as follows: Gapdh_F2(Gapdh5599): 5’-ACCACAGTCCATGCCATCAC-3’ Gapdh_R2(Gapdh5600): 5’-TCCACCACCCTGTTGCTGTA-3’ Zfp541_rtF2: 5’-CTGTTACTCAGAGGACCCCAGAAG-3’ Zfp541_rtR1: 5’-ATCTTTAACTCAGGAGTTGGACGG-3’ KCTD19_rtL1: 5’-GAAGACAATCTGCTGTGGCTG-3’ KCTD19_rtR1: 5’-GGGAACAGCCACCCATATTCA-3’ Primer sequences are listed in Table S3.

#### Preparation of testis extracts and immunoprecipitation

To prepare testis extracts, testes were removed from male C57BL/6 mice (P18-23), detunicated, and then resuspended in extraction buffer (20 mM Tris-HCl [pH 7.5], 100 mM KCl, 0.4 mM EDTA, 0.1% TritonX100, 10% glycerol, 1 mM β-mercaptoethanol) supplemented with Complete Protease Inhibitor (Roche). After homogenization, the cell extracts were filtrated to remove debris. The soluble chromatin-unbound fraction was collected after ultra-centrifugation at 100,000*g* for 30 min. The insoluble pellet was washed with buffer (10 mM Tris-HCl [pH 7.5], 1 mM CaCl_2_, 1.5 mM MgCl_2_, 10% glycerol) and digested with Micrococcus nuclease (0.008 units/ml) at 4°C for 60 min. The solubilized fractions were removed after centrifugation at 20,000*g* for 10 min at 4°C. The chromatin-bound fractions were extracted from the insoluble pellet by high salt extraction buffer (20 mM HEPES-KOH [pH 7.0], 400 mM KCl, 5 mM MgCl_2_, 0.1% Tween20, 10% glycerol, 1 mM β-mercaptoethanol) supplemented with Complete Protease Inhibitor. The solubilized chromatin fractions were collected after centrifugation at 100,000*g* for 30 min at 4°C.

For immunoprecipitation of endogenous ZFP541 and KCTD19 from extracts, 5 μg of affinity-purified rabbit anti-ZFP541, KCTD19-N and control IgG antibodies were crosslinked to 50 μl of protein A-Dynabeads (Thermo-Fisher) by DMP (Sigma). The antibody-crosslinked beads were added to the testis extracts prepared from WT testes (P13-18). The beads were washed with low salt extraction buffer. The bead-bound proteins were eluted with 40 μl of elution buffer (100 mM Glycine-HCl [pH 2.5], 150 mM NaCl), and then neutralized with 4 μl of 1 M Tris-HCl [pH 8.0]. The immunoprecipitated proteins were run on 4-12 % NuPAGE (Thermo-Fisher) in MOPS-SDS buffer and immunoblotted. Immunoblot image was developed using ECL prime (GE healthcare) and captured by LAS4000 (GE healthcare) or X-ray film.

#### Mass spectrometry

For mass spectrometry analysis of ZFP541 associated factors, fractions containing the ZFP541 immunoprecipitates were concentrated by precipitation with 10% trichloroacetic acid. The derived precipitates were dissolved in 7 M Urea, 50 mM Tris-HCl (pH 8.0), 5 mM EDTA solution, with 5 mM DTT at 37◻ for 30 min, and cysteine SH groups were alkylated with 10 mM iodoacetamide at 37◻ for 1 h. After alkylation, the solutions were desalted by methanol/chloroform precipitation, and the precipitates were dissolved in 2 M urea, 50 mM Tris-HCl buffer and subjected to trypsin gold (Promega) digestion overnight at 37◻. The resulting mixture of peptides was applied directly to the LC-MS/MS analysis system (Zaplous, AMR, Tokyo, Japan) using Finnigan LTQ mass spectrometry (Thermo Scientific) and a reverse phase C18 ESI column (0.2 × 50 mm, LC assist). The protein annotation data were verified in the mouse NCBI sequences using Bioworks software (Ver. 3.3; Thermo Scientific) with quantitation featuring the SEQUEST search algorithm. Two independent samples of ZFP541-IP from chromatin bound-, chromatin unbound-, MNase solubilized nucleosome fractions were analyzed. Identified proteins were presented after excluding the proteins detected in the control IgG-IP.

For mass spectrometry analysis of KCTD19-IP, the immunoprecipitated proteins were run on 4-12% NuPAGE (Thermo Fisher Scientific) by 1 cm from the well and stained with SimplyBlue (LC6065, Thermo Fisher Scientific) for the in-gel digestion. The gel that contained proteins was excised, cut into approximately 1mm sized pieces. Proteins in the gel pieces were reduced with DTT (20291, Thermo Fisher Scientific), alkylated with iodoacetamide (90034, Thermo Fisher Scientific), and digested with trypsin and lysyl endopeptidase (Promega, USA) in a buffer containing 40 mM ammonium bicarbonate, pH 8.0, overnight at 37 °C. The resultant peptides were analyzed on an Advance UHPLC system (AMR/Michrom Bioscience) coupled to a Q Exactive mass spectrometer (Thermo Fisher Scientific) processing the raw mass spectrum using Xcalibur (Thermo Fisher Scientific). The raw LC-MS/MS data was analyzed against the NCBI non-redundant protein and UniProt restricted to *Mus musculus* using Proteome Discoverer version 1.4 (Thermo Fisher Scientific) with the Mascot search engine version 2.5 (Matrix Science). A decoy database comprised of either randomized or reversed sequences in the target database was used for false discovery rate (FDR) estimation, and Percolator algorithm was used to evaluate false positives. Search results were filtered against 1% global FDR for high confidence level. Identified proteins were presented after excluding the proteins detected in the control IgG-IP.

#### Antibodies

The following antibodies were used for immunoblot (IB) and immunofluorescence (IF) studies: rabbit anti-Actin (IB, 1:2000, sigma A2066), rabbit anti-SYCP1 (IF, 1:1000, Abcam ab15090), mouse anti-γH2AX (IF, 1:1000, Abcam ab26350), rabbit anti-γH2AX (IF, 1:1000, Abcam ab11174), mouse anti-SYCP3 (Ishiguro et al., 2011), rabbit anti-MEIKIN (Kim et al., 2015) (IF, 1:1000), rabbit anti-H3S10P (IF, 1:2000, ab5176), rabbit anti-TDIF1 (IB, 1:1000, ab228703), rabbit anti-HDAC2 (IB, 1:1000, Abcam ab32117), mouse anti-HDAC1 (1:1000, Upstate 05-614), rabbit TDIF1 (IB, 1:1000, Abcam ab228703), mouse anti-SYCP1 (IF, 1:1000) (Ishiguro et al., 2011), rat anti-SYCP3 (Ishiguro et al., 2020) (IF, 1:1000), gunia pig anti-SYCP3 (Ishiguro et al., 2020) (IF, 1:2000), guinea pig anti-H1t (IF, 1:2000, kindly provided by Marry Ann Handel) (Cobb et al., 1999), α-tubulin DM1A (IB, 1:2000, Sigma), rabbit anti-DAZL (IF, 1:1000, ab34139).

His-tagged recombinant proteins of ZFP541-N (aa 1-517), KCTD19-N (aa1-308) and KCTD19-C (aa 694-950) were produced by inserting cDNA fragments in-frame with pET28c (Novagen) in *E. coli* strain BL21-CodonPlus(DE3)-RIPL (Agilent), solubilized in a denaturing buffer (6 M HCl-Guanidine, 20 mM Tris-HCl [pH 7.5]) and purified by Ni-NTA (QIAGEN) under denaturing conditions. Polyclonal antibodies against ZFP541-N were generated by immunizing rabbits. Polyclonal antibodies against KCTD19-N were generated by immunizing rabbits, rats and mice. Polyclonal antibodies against mouse KCTD19-C were generated by immunizing rabbits and rats. The antibodies were affinity-purified from the immunized serum with immobilized antigen peptides on CNBr-activated Sepharose (GE healthcare).

#### Histological Analysis

For hematoxylin and eosin staining, testes, epididymis and ovaries were fixed in 10% formalin or Bouin solution, and embedded in paraffin. Sections were prepared on MAS-GP typeA-coated slides (Matsunami) at 6 μm thickness. The slides were dehydrated and stained with hematoxylin and eosin.

For Immunofluorescence staining, testes and embryonic ovaries were embedded in Tissue-Tek O.C.T. compound (Sakura Finetek) and frozen. Cryosections were prepared on the MAS-GP typeA-coated slides (Matsunami) at 8 μm thickness, and then air-dried and fixed in 4% paraformaldehyde in PBS at pH◻7.4. The serial sections of frozen testes were fixed in 4% PFA for 5 min at room temperature and permeabilized in 0.1% TritonX100 in PBS for 10 min. The sections were blocked in 3% BSA/PBS or Blocking One (Nakarai), and incubated at room temperature with the primary antibodies in a blocking solution. After three washes in PBS, the sections were incubated for 1 h at room temperature with Alexa-dye-conjugated secondary antibodies (1:1000; Invitrogen) in a blocking solution. TUNEL assay was performed using MEBSTAIN Apoptosis TUNEL Kit Direct (MBL 8445). DNA was counterstained with Vectashield mounting medium containing DAPI (Vector Laboratory).

#### Immunostaining of spermatocytes

Spread nuclei from spermatocytes were prepared as described previously (Ishiguro et al., 2014). Briefly testicular cells were suspended in PBS, then dropped onto a slide glass together with an equal volume of 2% PFA, 0.2% (v/v) Triton X-100 in PBS, and incubated at room temperature in humidified chamber. The sides were then air-dried and washed with PBS containing 0.1% Triton-X100 or frozen for longer storage at −80°C. The serial sections of frozen testes were fixed in 4% PFA for 5 min at room temperature and permeabilized in 0.1% TritonX100 in PBS for 10 min. The slides were blocked in 3% BSA/PBS, and incubated at room temperature with the primary antibodies in a blocking solution. After three washes in PBS, the sections were incubated for 1 h at room temperature with Alexa-dye-conjugated secondary antibodies (1:1000; Invitrogen) in a blocking solution. DNA was counterstained with Vectashield mounting medium containing DAPI (Vector Laboratory).

#### Imaging

Immunostaining images were captured with DeltaVision (GE Healthcare). The projection of the images were processed with the SoftWorx software program (GE Healthcare). All images shown were Z-stacked. Bright field images were captured with OLYMPUS BX53 fluorescence microscope and processed with CellSens standard program.

#### Giemsa staining of metaphase I chromosome spread

The spermatocytes were treated in hypotonic buffer for 10 min and fixed in the Carnoy’s Fixative (75 % Methanol, 25% Acetic Acid) and, stained in 3% Giemsa solution for 30min.

#### Chromatin Immunoprecipitation

The seminiferous tubules from male C57BL/6 mice (P18-19-old age) were minced and digested with accutase and 0.5 units/ml DNase II, followed by filtration through a 40 μm cell strainer (FALCON). Testicular cells were fixed in 1% formaldehyde (Thermo-Fisher)/2mM Disuccinimidyl glutarate (ProteoChem) in PBS for 10 minutes at room temperature. Crosslinked cells were lysed with LB1 (50 mM HEPES pH 7.5, 140 mM NaCl, 1 mM EDTA, 10% glycerol, 0.5% NP-40, and 0.25% Triton X-100) and washed with LB2 (10 mM Tris-HCl pH 8.0, 200 mM NaCl, 1 mM EDTA, 0.5 mM EGTA). Chromatin lysates were prepared in LB3 (50 mM Tris-HCl pH8.0, 1% SDS, 10 mM EDTA, proteinase inhibitor cocktail (Sigma)), by sonication with Covaris S220 (Peak Incident Power, 175; Acoustic Duty Factor, 10%; Cycle Per Burst, 200; Treatment time, 600sec; Cycle, 2).

ZFP541 ChIP was performed using chromatin lysates and protein A Dyna-beads (Thermo-Fisher) coupled with 5 μg of rabbit anti-ZFP541-N antibodies (#1 and #2). After 4 hours of incubation at 4 °C, beads were washed 4 times in a low salt buffer (20 mM Tris-HCl (pH 8.0), 0.1% SDS, 1% (w/v) TritonX-100, 2 mM EDTA, 150 mM NaCl), and two times with a high salt buffer (20 mM Tris-HCl (pH 8.0), 0.1% SDS, 1% (w/v) TritonX-100, 2 mM EDTA, 500 mM NaCl). Chromatin complexes were eluted from the beads by agitation in elution buffer (10 mM Tris-HCl (pH 8.0), 300 mM NaCl, 5 mM EDTA, 1% SDS) and incubated overnight at 65 °C for reverse-crosslinking. Eluates were treated with RNase A and Proteinase K, and DNA was ethanol precipitated.

#### RNA-seq Data Analysis

GFP positive cells from *Rec8*-*3xFLAG-HA-p2A-GFP* knock-in male testis were sorted using SH800S cell sorter (SONY). Total RNAs were prepared by Trizol (Thermo Fisher) and quality of total RNA was confirmed by BioAnalyzer 2100 (RIN > 8) (Agilent). Library DNAs were prepared according to the Illumina Truseq protocol using Truseq Standard mRNA LT Sample Prep Kit (Illumina) and sequenced by Illumina NextSeq 500 (Illumina) using Nextseq 500/550 High Output v2.5 Kit (Illumina) to obtain single end 75 nt reads.

Resulting reads were aligned to the mouse genome UCSC mm10 using STAR ver.2.6.0a after trimmed to remove adapter sequence and low-quality ends using Trim Galore! v0.5.0 (cutadapt v1.16). Gene expression level measured as RPKM was determined by Cuffdiff v 2.2.1. Differential expression analysis using TPM was done by RSEM v1.3.1. GTF file was derived from UCSC mm10. Principal component analysis and GO-term analysis were performed using DAVID Bioinformatics Resources 6.8 (Huang da et al., 2009).

#### ChIP-seq Data Analysis

ChIP-seq libraries were prepared using 20 ng of input DNA, and 4 ng of ZFP541-ChIP DNA with KAPA Library Preparation Kit (KAPA Biosystems) and NimbleGen SeqCap Adaptor Kit A or B (Roche) and sequenced by Illumina Hiseq 1500 to obtain single end 50 nt reads.

ChIP-seq reads were trimmed to remove adapter sequence when converting to a fastq file. The trimmed ChIP-seq reads were mapped to the UCSC mm10 genome assemblies using Bowtie2 v2.3.4.1 with default parameters. Peak calling was performed using MACS program v2.1.0 (Zhang et al., 2008) (https://github.com/macs3-project/MACS) with the option (−g mm −p 0.00001). To calculate the number of overlapping peaks between Ab-1 and Ab-2, we used bedtools program (v2.27.1) (Quinlan and Hall, 2010) (https://bedtools.readthedocs.io/en/latest/). ChIP binding regions were annotated with BED file using Cis-regulatory Element System (CEAS) v0.9.9.7 (package version 1.0.2), in which the gene annotation table was derived from UCSC mm10. Motif identification was performed using MEME-ChIP v5.1.1 website (http://meme-suite.org/tools/meme-chip) (Bailey, 2011). The motif database chosen was “JASPAR Vertebrates and UniPROBE Mouse”. BigWig files, which indicate occupancy of ZFP541, were generated using deepTools (v3.1.0) and visualized with Integrative Genomics Viewer software (v.2.8.3) http://software.broadinstitute.org/software/igv/home. GO-term analyses were performed using DAVID Bioinformatics Resources 6.8 (Huang da et al., 2009) (https://david.ncifcrf.gov/). Aggregation plots and heatmaps for each sample against the dataset of ZFP541 target genes were made using deeptools program v3.5.0. The nearest genes of the ChIP-seq peaks were determined by GREAT website (v4.0.4) (http://great.stanford.edu/public/html/). Enrichment analyses of H3K27me3 and H3K27ac over ZFP541-bound targets were performed with bedtools program (v2.27.1) with −c option.

## DATA AND CODE AVAILABILITY

All data supporting the conclusions are present in the paper and the supplementary materials. Mouse lines generated in this study have been deposited to Center for Animal Resources and Development (CARD), *Zfp541*mutant mouse (ID 2857), *Kctd19* mutant Line #12 (ID 2879), *Kctd19* mutant Line #26 (ID 2880) and *Rec8*-*3xFLAG-HA-p2A-GFP* knock-in mouse (ID2681). The ZFP541 ChIP-seq data of mouse testes are deposited in the DDBJ Sequence Read Archive (DRA) under accession number GSE163916. RNA-seq data is deposited under GSE163917.

ChIP-seq data of H3K27me3 (Maezawa et al., 2018b) and H3K27ac (Maezawa et al., 2020) were derived from GSE89502 and GSE130652, respectively. Data for RNA-seq (THY1+SG, PS, RS) (Hasegawa et al., 2015) and RNA-seq (KIT+SG) (Maezawa et al., 2018b) were derived from GSE55060 and GSE89502, respectively.

## Reference

Alavattam, K.G., Maezawa, S., Sakashita, A., Khoury, H., Barski, A., Kaplan, N., and Namekawa, S.H. (2019). Attenuated chromatin compartmentalization in meiosis and its maturation in sperm development. Nat Struct Mol Biol 26, 175–184.

Bailey, T.L. (2011). DREME: motif discovery in transcription factor ChIP-seq data. Bioinformatics 27, 1653–1659.

Bantscheff, M., Hopf, C., Savitski, M.M., Dittmann, A., Grandi, P., Michon, A.M., Schlegl, J., Abraham, Y., Becher, I., Bergamini, G., et al. (2011). Chemoproteomics profiling of HDAC inhibitors reveals selective targeting of HDAC complexes. Nat Biotechnol 29, 255–265.

Burgoyne, P.S., Mahadevaiah, S.K., and Turner, J.M. (2009). The consequences of asynapsis for mammalian meiosis. Nat Rev Genet 10, 207–216.

Choi, E., Han, C., Park, I., Lee, B., Jin, S., Choi, H., Kim, D.H., Park, Z.Y., Eddy, E.M., and Cho, C. (2008). A novel germ cell-specific protein, SHIP1, forms a complex with chromatin remodeling activity during spermatogenesis. J Biol Chem 283, 35283–35294.

Chu, S., and Herskowitz, I. (1998). Gametogenesis in yeast is regulated by a transcriptional cascade dependent on Ndt80. Mol Cell 1, 685–696.

Cobb, J., Cargile, B., and Handel, M.A. (1999). Acquisition of competence to condense metaphase I chromosomes during spermatogenesis. Dev Biol 205, 49–64.

da Cruz, I., Rodriguez-Casuriaga, R., Santinaque, F.F., Farias, J., Curti, G., Capoano, C.A., Folle, G.A., Benavente, R., Sotelo-Silveira, J.R., and Geisinger, A. (2016). Transcriptome analysis of highly purified mouse spermatogenic cell populations: gene expression signatures switch from meiotic-to postmeiotic-related processes at pachytene stage. BMC Genomics 17, 294.

Drabent, B., Bode, C., Bramlage, B., and Doenecke, D. (1996). Expression of the mouse testicular histone gene H1t during spermatogenesis. Histochem Cell Biol 106, 247–251.

Ernst, C., Eling, N., Martinez-Jimenez, C.P., Marioni, J.C., and Odom, D.T. (2019). Staged developmental mapping and X chromosome transcriptional dynamics during mouse spermatogenesis. Nat Commun 10, 1251.

Handel, M.A., and Schimenti, J.C. (2010). Genetics of mammalian meiosis: regulation, dynamics and impact on fertility. Nat Rev Genet 11, 124–136.

Hasegawa, K., Sin, H.S., Maezawa, S., Broering, T.J., Kartashov, A.V., Alavattam, K.G., Ichijima, Y., Zhang, F., Bacon, W.C., Greis, K.D., et al. (2015). SCML2 establishes the male germline epigenome through regulation of histone H2A ubiquitination. Dev Cell 32, 574–588.

Hein, M.Y., Hubner, N.C., Poser, I., Cox, J., Nagaraj, N., Toyoda, Y., Gak, I.A., Weisswange, I., Mansfeld, J., Buchholz, F., et al. (2015). A human interactome in three quantitative dimensions organized by stoichiometries and abundances. Cell 163, 712–723.

Huang da, W., Sherman, B.T., and Lempicki, R.A. (2009). Systematic and integrative analysis of large gene lists using DAVID bioinformatics resources. Nat Protoc 4, 44–57.

Ichijima, Y., Sin, H.S., and Namekawa, S.H. (2012). Sex chromosome inactivation in germ cells: emerging roles of DNA damage response pathways. Cell Mol Life Sci 69, 2559–2572.

Ishiguro, K., Kim, J., Fujiyama-Nakamura, S., Kato, S., and Watanabe, Y. (2011). A new meiosis-specific cohesin complex implicated in the cohesin code for homologous pairing. EMBO Rep 12, 267–275.

Ishiguro, K., Kim, J., Shibuya, H., Hernandez-Hernandez, A., Suzuki, A., Fukagawa, T., Shioi, G., Kiyonari, H., Li, X.C., Schimenti, J., et al. (2014). Meiosis-specific cohesin mediates homolog recognition in mouse spermatocytes. Genes Dev 28, 594–607.

Ishiguro, K.I., Matsuura, K., Tani, N., Takeda, N., Usuki, S., Yamane, M., Sugimoto, M., Fujimura, S., Hosokawa, M., Chuma, S., et al. (2020). MEIOSIN Directs the Switch from Mitosis to Meiosis in Mammalian Germ Cells. Dev Cell 52, 429–445 e410.

Kim, J., Ishiguro, K., Nambu, A., Akiyoshi, B., Yokobayashi, S., Kagami, A., Ishiguro, T., Pendas, A.M., Takeda, N., Sakakibara, Y., et al. (2015). Meikin is a conserved regulator of meiosis-I-specific kinetochore function. Nature 517, 466–471.

Kimmins, S., and Sassone-Corsi, P. (2005). Chromatin remodelling and epigenetic features of germ cells. Nature 434, 583–589.

Kojima, M.L., de Rooij, D.G., and Page, D.C. (2019). Amplification of a broad transcriptional program by a common factor triggers the meiotic cell cycle in mice. Elife 8.

Kota, S.K., and Feil, R. (2010). Epigenetic transitions in germ cell development and meiosis. Dev Cell 19, 675–686.

Maezawa, S., Hasegawa, K., Alavattam, K.G., Funakoshi, M., Sato, T., Barski, A., and Namekawa, S.H. (2018a). SCML2 promotes heterochromatin organization in late spermatogenesis. J Cell Sci 131.

Maezawa, S., Hasegawa, K., Yukawa, M., Kubo, N., Sakashita, A., Alavattam, K.G., Sin, H.S., Kartashov, A.V., Sasaki, H., Barski, A., et al. (2018b). Polycomb protein SCML2 facilitates H3K27me3 to establish bivalent domains in the male germline. Proc Natl Acad Sci U S A 115, 4957–4962.

Maezawa, S., Sakashita, A., Yukawa, M., Chen, X., Takahashi, K., Alavattam, K.G., Nakata, I., Weirauch, M.T., Barski, A., and Namekawa, S.H. (2020). Super-enhancer switching drives a burst in gene expression at the mitosis-to-meiosis transition. Nat Struct Mol Biol 27, 978–988.

Namekawa, S.H., Park, P.J., Zhang, L.F., Shima, J.E., McCarrey, J.R., Griswold, M.D., and Lee, J.T. (2006). Postmeiotic sex chromatin in the male germline of mice. Curr Biol 16, 660–667.

Page, S.L., and Hawley, R.S. (2004). The genetics and molecular biology of the synaptonemal complex. Annu Rev Cell Dev Biol 20, 525–558.

Quinlan, A.R., and Hall, I.M. (2010). BEDTools: a flexible suite of utilities for comparing genomic features. Bioinformatics 26, 841–842.

Sasaki, H., and Matsui, Y. (2008). Epigenetic events in mammalian germ-cell development: reprogramming and beyond. Nat Rev Genet 9, 129–140.

Sawai, Y., Kasamatsu, A., Nakashima, D., Fushimi, K., Kasama, H., Iyoda, M., Kouzu, Y., Shiiba, M., Tanzawa, H., and Uzawa, K. (2018). Critical role of deoxynucleotidyl transferase terminal interacting protein 1 in oral cancer. Lab Invest 98, 980–988.

Schultz, N., Hamra, F.K., and Garbers, D.L. (2003). A multitude of genes expressed solely in meiotic or postmeiotic spermatogenic cells offers a myriad of contraceptive targets. Proc Natl Acad Sci U S A 100, 12201–12206.

Shima, J.E., McLean, D.J., McCarrey, J.R., and Griswold, M.D. (2004). The murine testicular transcriptome: characterizing gene expression in the testis during the progression of spermatogenesis. Biol Reprod 71, 319–330.

Sin, H.S., Kartashov, A.V., Hasegawa, K., Barski, A., and Namekawa, S.H. (2015). Poised chromatin and bivalent domains facilitate the mitosis-to-meiosis transition in the male germline. BMC Biol 13, 53.

Turnbull, R.E., Fairall, L., Saleh, A., Kelsall, E., Morris, K.L., Ragan, T.J., Savva, C.G., Chandru, A., Millard, C.J., Makarova, O.V., et al. (2020). The MiDAC histone deacetylase complex is essential for embryonic development and has a unique multivalent structure. Nat Commun 11, 3252.

Turner, J.M. (2015). Meiotic Silencing in Mammals. Annu Rev Genet 49, 395–412.

Wang, Y., Wang, H., Zhang, Y., Du, Z., Si, W., Fan, S., Qin, D., Wang, M., Duan, Y., Li, L., et al. (2019). Reprogramming of Meiotic Chromatin Architecture during Spermatogenesis. Mol Cell 73, 547–561 e546.

Xu, L., Ajimura, M., Padmore, R., Klein, C., and Kleckner, N. (1995). NDT80, a meiosis-specific gene required for exit from pachytene in Saccharomyces cerevisiae. Mol Cell Biol 15, 6572–6581.

Yagi, T., Tokunaga, T., Furuta, Y., Nada, S., Yoshida, M., Tsukada, T., Saga, Y., Takeda, N., Ikawa, Y., and Aizawa, S. (1993). A novel ES cell line, TT2, with high germline-differentiating potency. Anal Biochem 214, 70–76.

Zhang, Y., Liu, T., Meyer, C.A., Eeckhoute, J., Johnson, D.S., Bernstein, B.E., Nusbaum, C., Myers, R.M., Brown, M., Li, W., et al. (2008). Model-based analysis of ChIP-Seq (MACS). Genome Biol 9, R137.

Zhang, Y., Wang, Z., Huang, Y., Ying, M., Wang, Y., Xiong, J., Liu, Q., Cao, F., Joshi, R., Liu, Y., et al. (2018). TdIF1: a putative oncogene in NSCLC tumor progression. Signal Transduct Target Ther 3, 28.

Zickler, D., and Kleckner, N. (2015). Recombination, Pairing, and Synapsis of Homologs during Meiosis. Cold Spring Harb Perspect Biol 7.

